# Assembly reactions of SARS-CoV-2 nucleocapsid protein with nucleic acid

**DOI:** 10.1101/2023.11.22.568361

**Authors:** Huaying Zhao, Abdullah M. Syed, Mir M. Khalid, Ai Nguyen, Alison Ciling, Di Wu, Wai-Ming Yau, Sanjana Srinivasan, Dominic Esposito, Jennifer A. Doudna, Grzegorz Piszczek, Melanie Ott, Peter Schuck

**Affiliations:** Laboratory of Dynamics of Macromolecular Assembly, National Institute of Biomedical Imaging and Bioengineering, National Institutes of Health, Bethesda, MD 20892; Gladstone Institutes, San Francisco, CA 94158; Innovative Genomics Institute, University of California, Berkeley, CA 94720; Biophysics Core Facility, National Heart, Lung, and Blood Institute, National Institutes of Health, Bethesda, MD 20892; Laboratory of Chemical Physics, National Institute of Diabetes and Digestive and Kidney Diseases, National Institutes of Health, Bethesda, MD 20892; Protein Expression Laboratory, Frederick National Laboratory for Cancer Research, Frederick, MD 21702; Department of Molecular and Cell Biology, University of California, Berkeley, CA 94720; HHMI, University of California, Berkeley, CA 94720; Department of Chemistry, University of California, Berkeley, CA 94720; Molecular Biophysics and Integrated Bioimaging Division, Lawrence Berkeley National Laboratory, Berkeley, CA 94720; California Institute for Quantitative Biosciences (QB3), University of California, Berkeley, CA 94720; Department of Medicine, University of California, San Francisco, CA 94143; Chan Zuckerberg Biohub, San Francisco, CA 94158; Center for Biomedical Engineering Technology Acceleration, National Institute of Biomedical Imaging and Bioengineering, National Institutes of Health, Bethesda, MD 20892

## Abstract

The viral genome of SARS-CoV-2 is packaged by the nucleocapsid (N-) protein into ribonucleoprotein particles (RNPs), 38±10 of which are contained in each virion. Their architecture has remained unclear due to the pleomorphism of RNPs, the high flexibility of N-protein intrinsically disordered regions, and highly multivalent interactions between viral RNA and N-protein binding sites in both N-terminal (NTD) and C-terminal domain (CTD). Here we explore critical interaction motifs of RNPs by applying a combination of biophysical techniques to mutant proteins binding different nucleic acids in an *in vitro* assay for RNP formation, and by examining mutant proteins in a viral assembly assay. We find that nucleic acid-bound N-protein dimers oligomerize *via* a recently described protein-protein interface presented by a transient helix in its long disordered linker region between NTD and CTD. The resulting hexameric complexes are stabilized by multi-valent protein-nucleic acid interactions that establish crosslinks between dimeric subunits. Assemblies are stabilized by the dimeric CTD of N-protein offering more than one binding site for stem-loop RNA. Our study suggests a model for RNP assembly where N-protein scaffolding at high density on viral RNA is followed by cooperative multimerization through protein-protein interactions in the disordered linker.

## Introduction

SARS-CoV-2 is a persistent threat to global health and continued research is essential for the development of new vaccines and antivirals counteracting viral immune evasion and mutagenesis. Viral assembly is a critical step in its life cycle and potential target for therapeutic development, but so far only incompletely understood. As observed in recent cyro-electron tomography (cryo-ET) studies, virions contain ≈38±10 ribonucleoprotein particles (vRNPs) that condense the large viral genomic RNA (gRNA) like beads on a string (1, 2). vRNPs have 14 -16 nm sized pillar-shaped structures, are heterogeneous, and are composed of nucleocapsid (N-)protein and RNA with unknown architecture and assembly pathway.

N-protein is a 419 aa protein highly expressed in infected cells, and besides its key role in vRNP formation it is highly multifunctional (3–5). It consists in two folded domains, the NTD and CTD, which are linked and flanked by large intrinsically disordered regions (IDRs) termed N-arm, linker, and C-arm (6) (**Figure 1A**). The NTD forms a nucleic acid (NA) binding site that is thought to harbor recognition mechanism of a packaging signal (7). The CTD provides a high-affinity dimerization interface and also binds NA (6) (**Figure 1B**). Ultra-weak protein-protein interactions allow N-protein to undergo liquid-liquid phase separation (LLPS), enhanced in the presence of NA, to form droplets of high macromolecular density (8–11), which are thought to facilitate the assembly of vRNPs (9, 11–13). IDRs also play important roles in N-protein NA interactions (14–17). For example, the N-arm undergoes conformational changes upon NA binding to NTD, significantly increasing the affinity of the NTD for RNA (15). Similar to the N-arm, an SR-rich portion of the linker IDR flanking the NTD is also perturbed by NA binding (14), as is an L-rich sequence of the linker IDR (LRS) further downstream. Recently, we have shown that highly conserved transient helices in the LRS can assemble cooperatively into coiled-coil trimers, tetramers, and higher oligomers, in a process that is allosterically enhanced by NA binding (17, 18) (**Figure 1C**). We hypothesized that this mechanism might initiate assembly by providing additional protein-protein interfaces essential for stabilizing three-dimensional structures of the vRNPs (17, 18).

**Figure 1.**
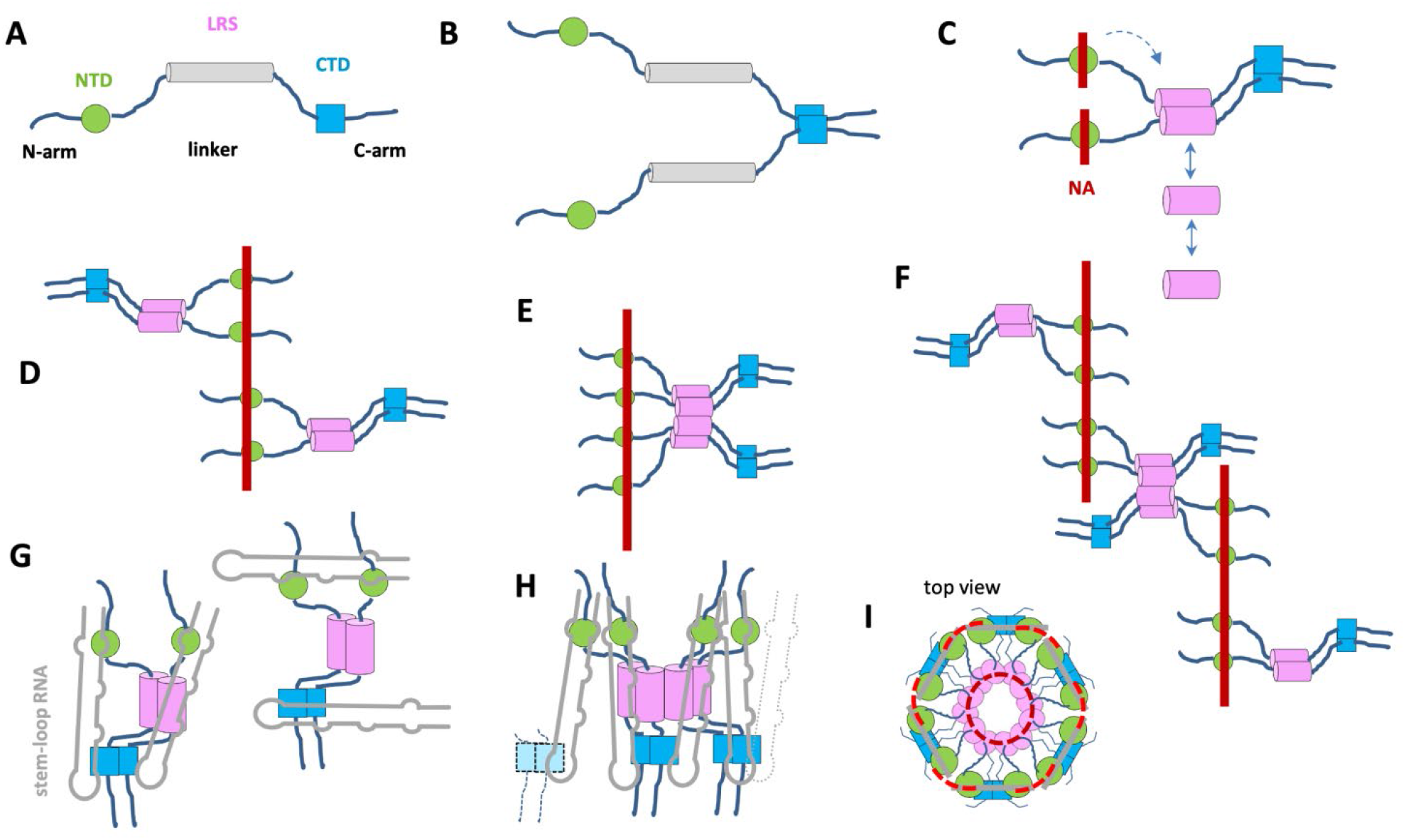
Schematic organization of N-protein and hypothetical architectures of NA complexes. (A) Organization of N-protein chain with folded domains NTD (green cylinder) and CTD (blue square), and IDRs N-arm, C-arm, and linker (cylinder), the latter containing the transient helix in the LRS capable of oligomerization. (B) N-protein in solution is a dimer linked with high affinity at the CTD. (C) The NA- ligated induction of folding in the LRS causes compaction (magenta) and LRS oligomerization in dimers, trimers, tetramers, and higher oligomers. (D) Configuration of two N-protein dimers independently scaffolded on NA T_40_ (red bar) without further LRS oligomerization. (E) Two N-protein dimers scaffolded on NA and stabilized through LRS oligomerization. (F) Similar to (E) with crosslink between NA strands allowing the formation of higher oligomers. (G) N-protein dimer with two SL7 stem-loop RNA ({2N/2SL7}, grey), depicted in alternate configurations occupying all major NA interfaces in the NTD and CTD creating different intra-dimer crosslinks. (H) Possible architecture of N-protein/stem-loop complexes allowing {2N/2SL7} units to oligomerize *via* LRS and simultaneous multi-valent binding of SL7 in inter-dimer crosslinks. Dotted lines belong to neighboring {2N/2SL7} units not fully drawn. (I) The top view of a 6x{2N/2SL7} hexamer of dimers. The dashed lines indicates two levels of inter-dimer crosslinks of neighboring {2N/2SL7} subunits, *via* contacts of the LRS interfaces (dark red dashed) and *via* multivalent RNA binding of CTD and/or scaffolding of NTD (light red dashed).

To study the assembly mechanism in more detail we examine N-protein binding to short NA probes of sufficient length for multi-valent binding and ask how scaffolding N-protein might diminish or augment simultaneous LRS oligomerization. As shown by Carlson and co-workers, mixtures of stem-loop RNA and N-protein form large RNA/protein complexes (RNPs) that may serve as a model for the viral vRNPs (11, 16). In the current study, to further elucidate the assembly mechanism we utilize this system and compare the size distribution and composition of complexes formed by wildtype N-protein (N_WT_) or mutants that abrogate LRS oligomerization binding either short linear oligonucleotides or stem-loop RNA. In parallel, we study the impact of the same mutations on viral packaging and assembly in a VLP assay. Our data highlight the role of the LRS oligomerization and the interplay with nucleic acid binding of NTD and CTD in the formation of RNPs. Taken together, these elements suggest features of possible RNP assembly models on the level of protein domains and their interfaces.

## Results

### LRS augments protein-NA complex formation in the presence of multi-valent NA binding

In previous work we have studied N-protein dimer in mixtures with sufficiently short oligonucleotides (T_10_) to eliminate the possibility of multi-valent binding, which allowed us to focus on the protein-protein interactions that arise for NA-liganded N-protein through allosteric stabilization of LRS helices and their cooperative coiled-coil oligomerization (17), as schematically indicated in **Figure 1C**. Occupation of the NA binding site (presumably in the NTD) causes a conformational change and improves the effective K_D_* for oligomerization by approximately three orders of magnitude from ≈1 mM to low µM. However, this is still significantly weaker than the N-protein affinity for NA. Therefore the question arises if the LRS interface can have a role in assembly when the N-protein dimer is more strongly and multivalently binding to NA. Possible scenarios for multivalent NA binding are sketched in **Figures 1D/E**, where either LRS oligomerization is not possible in N/NA complexes due to the strong NA binding interfaces constraining the complex structure (**Figure 1D**), or alternatively, complex architectures do permit LRS interactions between N-protein dimers while N-protein is scaffolded on NA (**Figure 1E/F**). The distinction is crucial for possible RNP assembly pathways.

To shed light on this question we studied N-protein binding to T_40_ oligonucleotides, which are twice the length required for cross-linking N-protein (19), and four times the size of the NTD binding epitope (20). As we have shown previously (17) it is conveniently possible to reduce or eliminate LRS oligomerization through LRS mutations that suppress (L222P) or completely disrupt (L222P/R226P) helix formation, rendering N-protein capable only of presenting protein/NA interfaces (besides the obligatory high-affinity dimerization of the CTD). Thus, as a starting point to probe the binding capacity of T_40_ for N-protein we carried out experiments in a concentration series with different molar ratios of mutant and wild-type (WT) N-protein binding to T_40_. Sedimentation velocity analytical ultracentrifugation (SV-AUC) permits size-dependent hydrodynamic separation of complexes and unbound species, and can be recorded in multiple optical signals to determine the molar ratio of protein and NA in the sedimenting macromolecular complexes (21, 22).

The resulting sedimentation coefficient distributions are shown in **Figure 2A**. In moderate ionic strength buffer B_150Na_ (20 mM HEPES, 150 mM NaCl, pH 7.4) in mixtures with tenfold molar excess of either N:L222P or N:L222P/R226P over T_40_ we observe a ≈6.7 S peak (dashed blue and cyan curves, respectively). This is distinctly higher than the sedimentation coefficient of ≈4 S of N-protein alone, and ≈1.7 S of free T_40_. It is also distinctly higher than complexes observed in analogous experiments with the shorter T_10_, that when bound to the same mutant protein produces only ≈5 S species consisting of T_10_- liganded N dimers (17). Unraveling the signal ratios of the 6.7 S N:L222P/T_40_ complex leads to a molar ratio of ≈4 N-proteins per T_40_, suggesting a highly extended scaffolded configuration such as sketched in **Figure 1D** (with an *s*-value of 6.7 S implying a translational frictional ratio f/f_0_ ≈ 2). At only threefold molar excess of these LRS mutants over T_40_ smaller complexes sedimenting at ≈5.2 S are observed (solid blue and cyan curves) with molar ratios protein/NA of ≈2:1, consistent with lower saturation of NA sites with a single N-protein dimer bound per T_40_.

**Figure 2:**
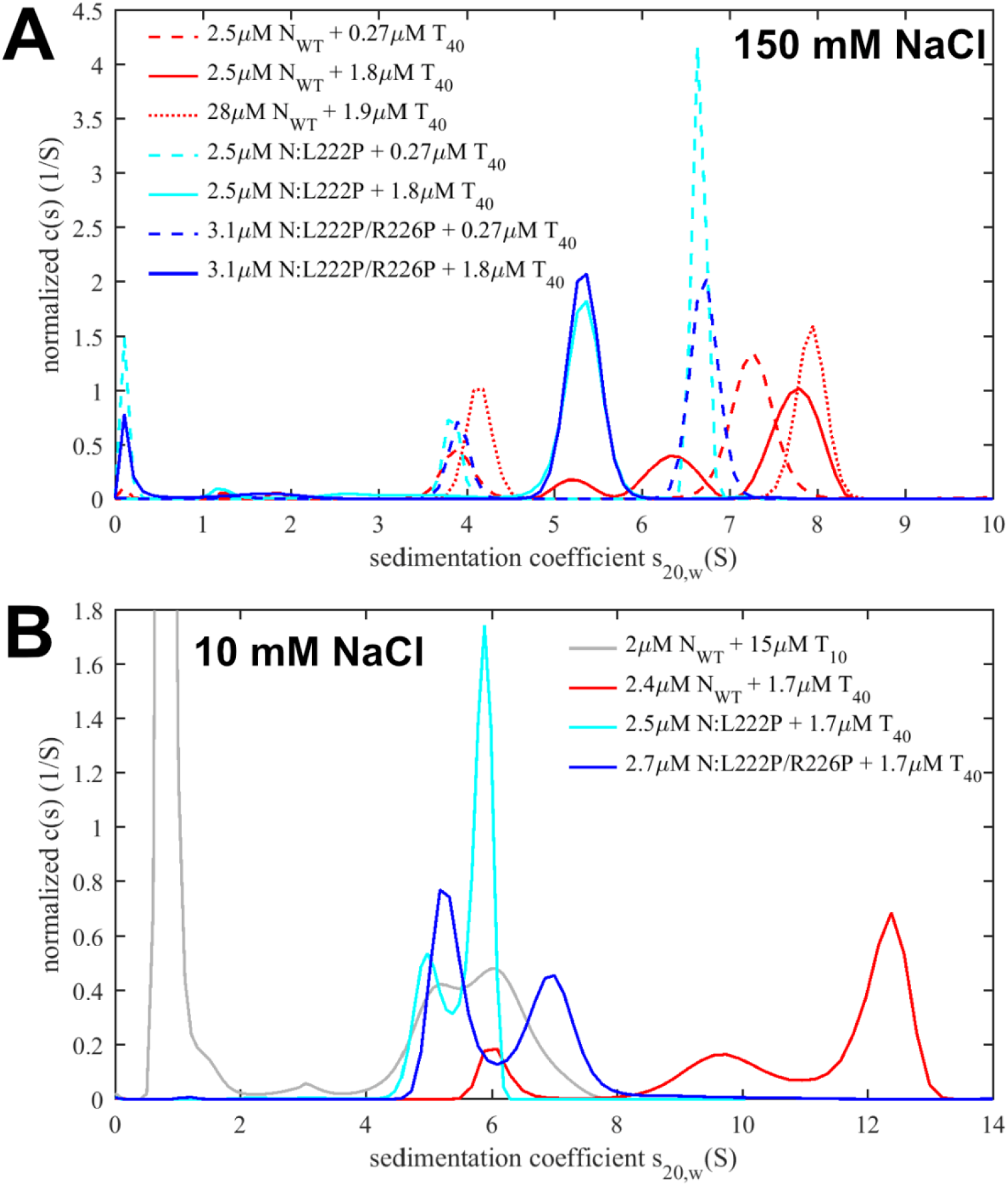
Simultaneous LRS oligomerization and scaffolding on a linear oligonucleotide T_40_. Shown are sedimentation coefficient distributions recorded at 260 nm for N_WT_ (red) and the LRS mutants N:L222P (cyan) and N:L222P/R226P (blue) at concentrations indicated. (A) Experiments in moderate ionic strength (PBS for 28 µM N_WT_ + 1.9 µM T_10_, all others buffer B_150Na_). (B) Complex formation at low ionic strength in buffer B_10Na_. For comparison, data with T_10_ are reproduced from (17).

By contrast, mixtures of N_WT_ with T_40_ at the same concentrations exhibit higher sedimentation coefficients throughout (**Figure 2A**, red curves). An increase in sedimentation coefficient with increasing total complex concentration can be discerned, as is characteristic in a ‘reaction boundary’, which signifies higher-order oligomerization rapidly reversible on the timescale of sedimentation (≈ 1,000 sec). The reaction boundaries reflect the time-average sedimentation velocity of all co-sedimenting complex and free species, and therefore represent a lower limit for the *s*-value of the largest complex (21, 23).

We can attribute this complex formation to the additional LRS protein-protein interfaces in N_WT_ augmenting complex formation, as in configuration **Figure 1F**. In the mixtures of 2.5 µM (or 28 µM) N_WT_ with ≈2 µM T_40_ a sedimentation boundary of ≈7.8 S is observed (**Figure 2A**, red solid line), as compared to the value of ≈5.2 S for the LRS mutants under the same conditions (blue and cyan solid lines). For reference, aside from complications in the interpretation of sedimentation velocities from reversible reacting systems (17), and potential conformational changes, such a 1.5-fold increase in sedimentation coefficient suggests the presence of majority species with at least twofold molecular weight . From the data at the highest concentration, a complex molar ratio of 4:1 N_WT_/T_40_ can be measured, which may correspond for example to complexes sketched in **Figure 1F**, although the detailed configuration is undetermined by the current data and may be heterogeneous.

SARS-CoV-2 N-protein is basic and can therefore be expected to exhibit strong electrostatic contributions to the driving force of NA binding and a strong dependence on ionic strength (24). We therefore examined the same N-protein/T_40_ interaction next at low salt buffer B_10Na_ (10 mM NaCl, 10.1 mM Na_2_PO_4_ , 1.8mM KH_2_PO_4_ , 2.7 mM KCl, pH 7.4) under otherwise identical conditions. Since previous studies quantifying the LRS oligomerization affinities were conducted with isolated LRS peptide (N:210- 246) at 150 mM NaCl (17), we carried out preliminary SV-AUC experiments with the LRS peptide in B_10Na_, and observed enhanced cooperative assembly of the LRS protein-protein interface in the lower ionic strength (**Supplementary Figure S1**). Thus, both the protein-NA interfaces and the protein-protein interfaces strengthen at lower ionic strength.

We examined the impact of this on complex formation. For the LRS mutants with ablated LRS interfaces at a 3:2 molar ratio of protein/T_40_ the ≈5 S species observed previously at moderate ionic strength is at low ionic strength accompanied by majority ≈6 S peak or minority ≈7 S peaks for N:L222P and N:L222P/R226P, respectively, indicating some additional assembly (or greater T_40_ saturation) (**Figure 2B**, blue and cyan curves). Low salt conditions also lead to larger complex formation with N_WT_-protein, but here exhibiting a significantly augmented effect: Under identical low salt conditions, N_WT_ forms complex species with T_40_ sedimenting at > 12.5 S (red curve). Based on the hydrodynamic scaling law *s* ∼ *M*^2/3^, the more than two-fold increase in sedimentation coefficient introduced by the LRS in N_WT_ compared to N:L222P implies a majority species with approximately threefold molecular weight. A back-of-the-envelope estimate suggests moderately compact (f/f_0_ = 1.3) species with *s*-values of 12.5 S require complexes of ≈300 kDa, and species of higher molecular weights if complexes are in more extended conformations or in reaction boundaries. This demonstrates the potential for strong oligomerization of LRS with complexes of even higher assembly state than sketched in **Figure 1E** and **F**, involving at least three N_WT_ dimers. It should be noted that in these low salt conditions, higher protein and T_40_ concentrations (such as 10 µM WT N-protein with 5 µM T_40_) promote the formation of macromolecular condensates.

SV-AUC data at intermediate ionic strength with majority potassium ions (buffer B_65K_ containing 64.8 mM KCl, 5.6 mM NaCl, 24.6 mM HEPES, pH 7.50) – a condition applied to the stem-loop RNP assembly assay described below – lead to results similar to those at the higher ionic strength B_150Na_ , reproducing the increased complex formation with T_40_ of N_WT_ as compared to both LRS mutants under otherwise identical conditions (**Supplementary Figure S2**).

Finally, we can observe the molar mass distribution directly using mass photometry (MP). This technique is applicable to mixtures at nanomolar concentrations, and is therefore suitable to explore the onset of binding. As shown in **Figure 3A**, in moderate ionic strength (buffer B_65K_) at 25 nM N_WT_ with 31 nM T_40_, the peak is indistinguishable from that of N_WT_ alone, which shows the expected dimeric state. Unbound T_40_ is below the minimum detectable mass in the MP instrument. At higher concentrations, increasingly the main peak shifts to slightly higher masses consistent with binding of T_40_, and secondary peak emerges with masses up to the ≈200 kDa range indicating onset of tetramerization at 0.25 µM N_WT_ with 0.31 µM T_40_. Consistent with this, SV-AUC of this mixture shows a reaction boundary at ≈6 S, at tenfold higher concentrations growing to ≈8 S (**Supplementary Figure S2**).

**Figure 3:**
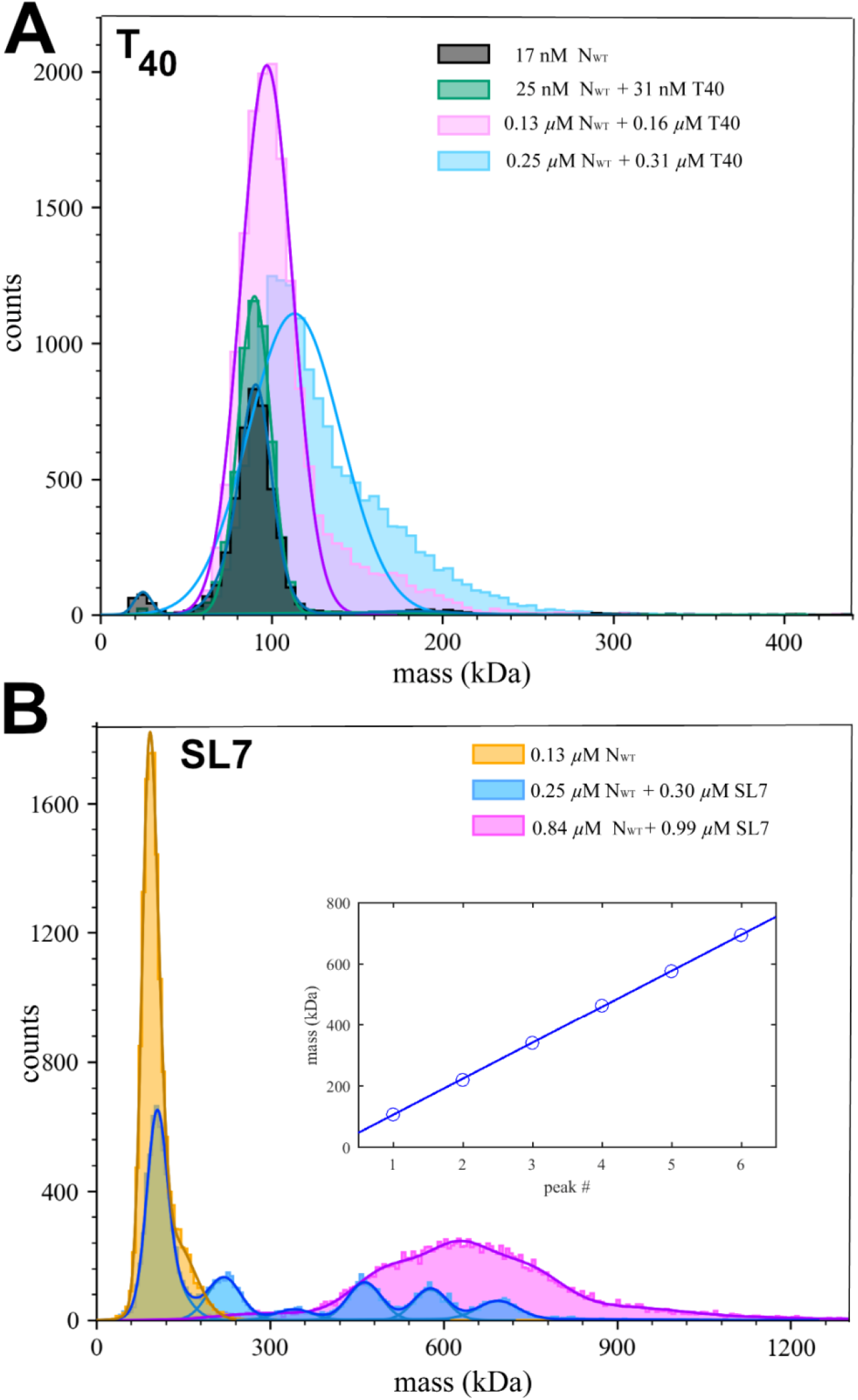
Complex mass distributions of N_WT_ with T_40_ and SL7 in mass photometry. (A) Concentration series of N_WT_ with T_40_; (B) Concentration series of N_WT_ with SL7. The inset shows peak mass values *vs* peak number of the 0.25 µM N_WT_ with 0.3 µM SL7 mixture and linear fit leading to a mass increment of 118 kDa.

### N-protein forms large RNP complexes with stem-loop RNA

Recently, the Morgan laboratory has examined binding of N-protein to viral genome segments of different length from the 5’ end of the viral genome, and observed the formation of ≈700-800 kDa complexes with 600 nt RNA consistent in size with vRNP particles observed in cryo-ET, and consistent in the average length of RNA that would need to be packaged by each vRNP (11, 16). Similar sized protein/RNA complexes were observed for N-protein binding to stem-loops SL4a, SL7, and SL8, which exhibited dissociation at submicromolar concentrations unless chemically crosslinked (16). In the present work, we focused on non-covalent binding of N-protein to stem-loop RNA SL7 (46 nt, 14.9 kDa) to compare it with the similar-sized DNA oligonucleotides T_40_.

Under the same conditions as shown in **Figure 3A** for T_40_, a mixture of 0.25 µM N_WT_ with 0.3 µM SL7 exhibits a distinct ladder of species with peak molar masses of 108 ± 24, 221 ± 29, 342 ± 33, 464 ± 29, 577 ± 29, 695 ± 90 kDa (**Figure 3B**, blue histogram), analogous to the results reported by Carlson et *al*. for SL8 (16). The best-fit increment is 118 kDa, which is closest to the theoretical molar mass of 121 kDa of an N_WT_ dimer in complex with two SL7. The first peak appears at a slightly lower mass, which may indicate a mixture of unresolved liganded and free N_WT_ dimer (free SL7 is below the mass limit of detection). Thus, the MP data are best consistent with an oligomerization of up to six {N_WT_ dimer/2SL7} subunits. However, we attribute an estimated ≈5-10% uncertainty to the molar mass values of MP due to differences in calibration, and differences in mass-to-contrast ratios for nucleic acids compared to proteins (25). (Solvent-dependent variation may be expected due to the strong counter-ion dependence of the refractive index of nucleic acids (26)). Therefore, complexes of N_WT_ dimer with one or three SL7 cannot be ruled out from the MP data.

As mentioned above, MP is limited to nanomolar macromolecular concentrations to allow discrimination of individual adsorption events. When the limit is approached at 0.84 µM N_WT_ + 0.99 µM SL7, as shown in **Figure 3B**, a single broad peak is observed ranging from 400 kDa to >900 kDa, suggesting the possibility of higher complexes at higher concentrations. Therefore, to study whether the complex formation with stem-loop RNA is an unlimited self-assembly or leads to a well-defined complex we turn back to hydrodynamic techniques.

Figure 4A shows a concentration series of N_WT_ and SL7 at constant molar ratio in SV-AUC. At low concentrations of 0.25 µM N_WT_ with 0.3 µM SL7 a broad distribution of species in the range of 7 – 18 S can be discerned (black line). This is qualitatively consistent with the ladder of complex species observed in MP under identical conditions (Figure 3B), although consideration is required of the fact that MP reports number distributions whereas SV-AUC produces weight-based distributions, i.e., the latter causes larger signals for larger particles. The ≈18 S peak confirms the strongly enhanced assembly with SL7 compared to T_40_. In Figure 4A, the 0.25 µM N_WT_ + 0.3 µM SL7 sample was created both by mixing of separate stocks to achieve the final concentration, or alternatively by 90 min incubation of a 10fold concentrated stock mixture followed by dilution. Within error the sedimentation coefficient distributions are indistinguishable, demonstrating they reflect equilibrium between species (after further 90 min incubation prior to the start of centrifugation).

**Figure 4:**
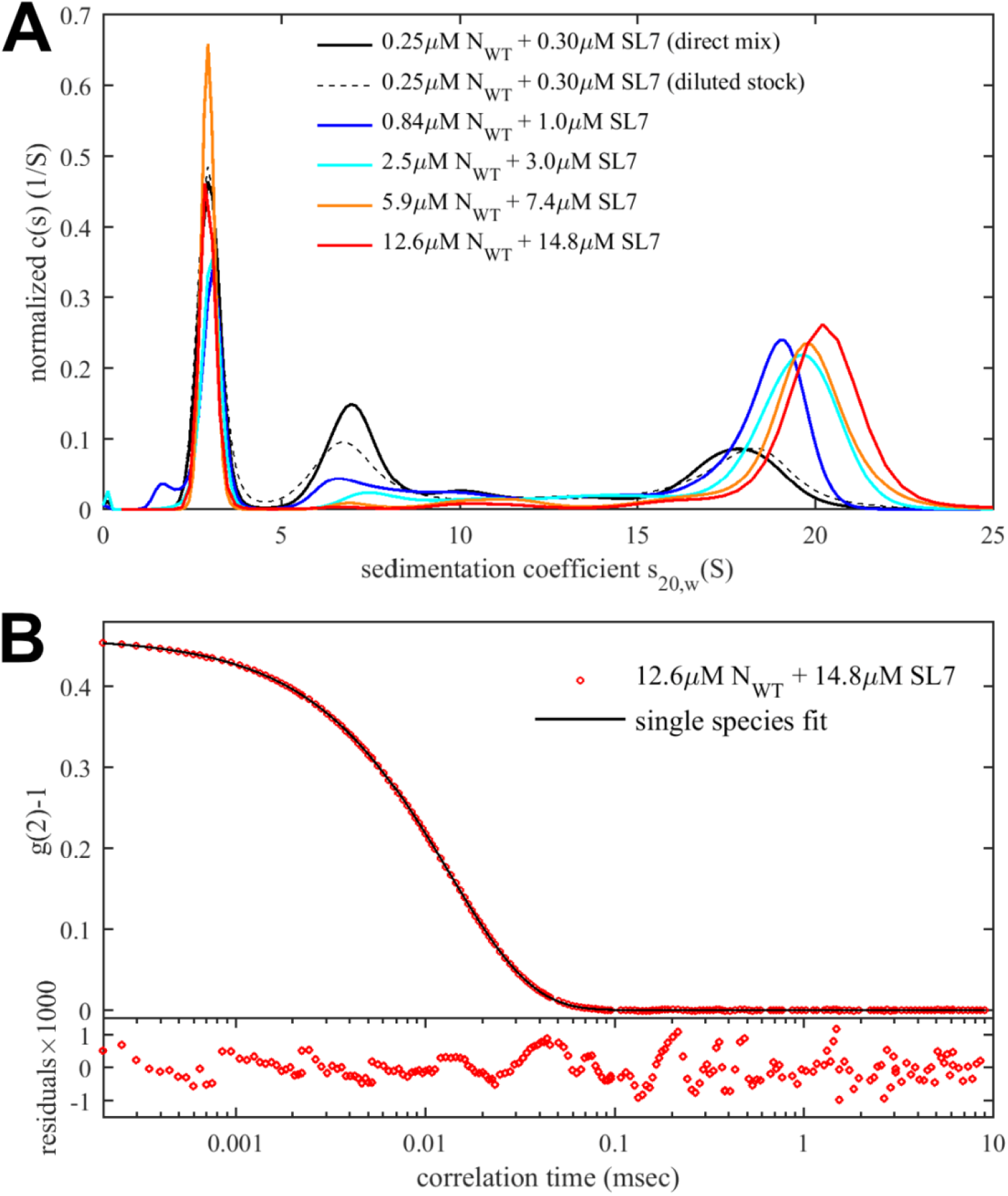
Concentration-dependent RNP assembly of N_WT_ with stem-loop SL7. (A) Sedimentation coefficient distributions from SV-AUC experiments recorded at 260 nm. Mixtures of N_WT_ and SL7 at concentrations indicated in B_65K_ (or B_75Na_ for the two most concentrated mixtures). (B) Autocorrelation data from DLS of the highest concentration mixture. The best-fit single-species model leads to a diffusion coefficient of 2.466 × 10^-7^ cm^2^/sec or a Stokes radius of 8.5 nm.

Increasing concentrations up to 50-fold (12.6 µM N_WT_ with 14.8 µM SL7) causes the largest species to asymptotically approach a sedimentation coefficient of ≈20 S. Multi-signal analysis of the fastest sedimenting boundary leads to a molar ratio of (0.95 ± 0.1) SL7/N_WT_, consistent with a model of self- assembly of {N_WT_ dimer/2SL7} subunits. In order to determine the mass of the largest RNP complex, we carried out DLS measurements. Even though this technique does not resolve species of similar size, the scattering intensity is weighted strongly in favor of the largest component. As shown in Figure 4B, the autocorrelation function can be fit well with a single species model with a diffusion coefficient of 2.45 F, corresponding to a Stokes radius of 8.5 nm. The diffusion coefficient can be combined with the peak sedimentation coefficient in the Svedberg equation *M* (1− *v ρ* ) = *RT* (*s D*) to produce a buoyant molar mass *M* (1− *v ρ* ) of ≈200 kDa. With a 1:1 molar ratio we estimate the complex partial-specific volume to be 0.695 mL/g, leading to a complex molar mass of ≈650 kDa. This estimate is slightly below the value from MP of ≈695 kDa, as well as the theoretical value of 726 kDa for a hexamer {N_WT_ dimer/2SL7}_6_, which may be caused by an underestimate of the largest species *s*-value from inspection of the reaction boundaries with incomplete saturation (21) and a overestimate of the diffusion coefficient due to the unaccounted scattering contributions of free SL7. Nevertheless, it clearly shows the finite assembly of N_WT_ with SL7 to species not exceeding a hexamer of {N_WT_ dimer/2SL7} subunits. Finally, from the molar mass and sedimentation coefficient we can calculate the frictional ratio and arrive at a value of 1.5; this indicates the complex is significantly more compact than the free N-protein which has a frictional ratio of 1.8 (19).

### The formation of large RNP complexes depends on LRS oligomerization and multivalent NA binding

Analogous to our studies of N-protein binding to T_40_ above, we next studied how the LRS protein-protein interfaces contribute to the formation of RNP complexes by examining the reaction products of SL7 with LRS point mutants N:L222P and N:L222P/R226P. First, Figure 5A shows the mass distribution obtained at low concentrations suitable for MP experiments (0.25 µM N-protein with 0.3 µM SL7). While for N_WT_ in this mixture a ladder of complex species up to ≈700 kDa can be discerned, very similar to the experiment shown in Figure 3B (of which it is a replicate), both LRS mutants show a single peak consistent with N-protein dimer.

**Figure 5:**
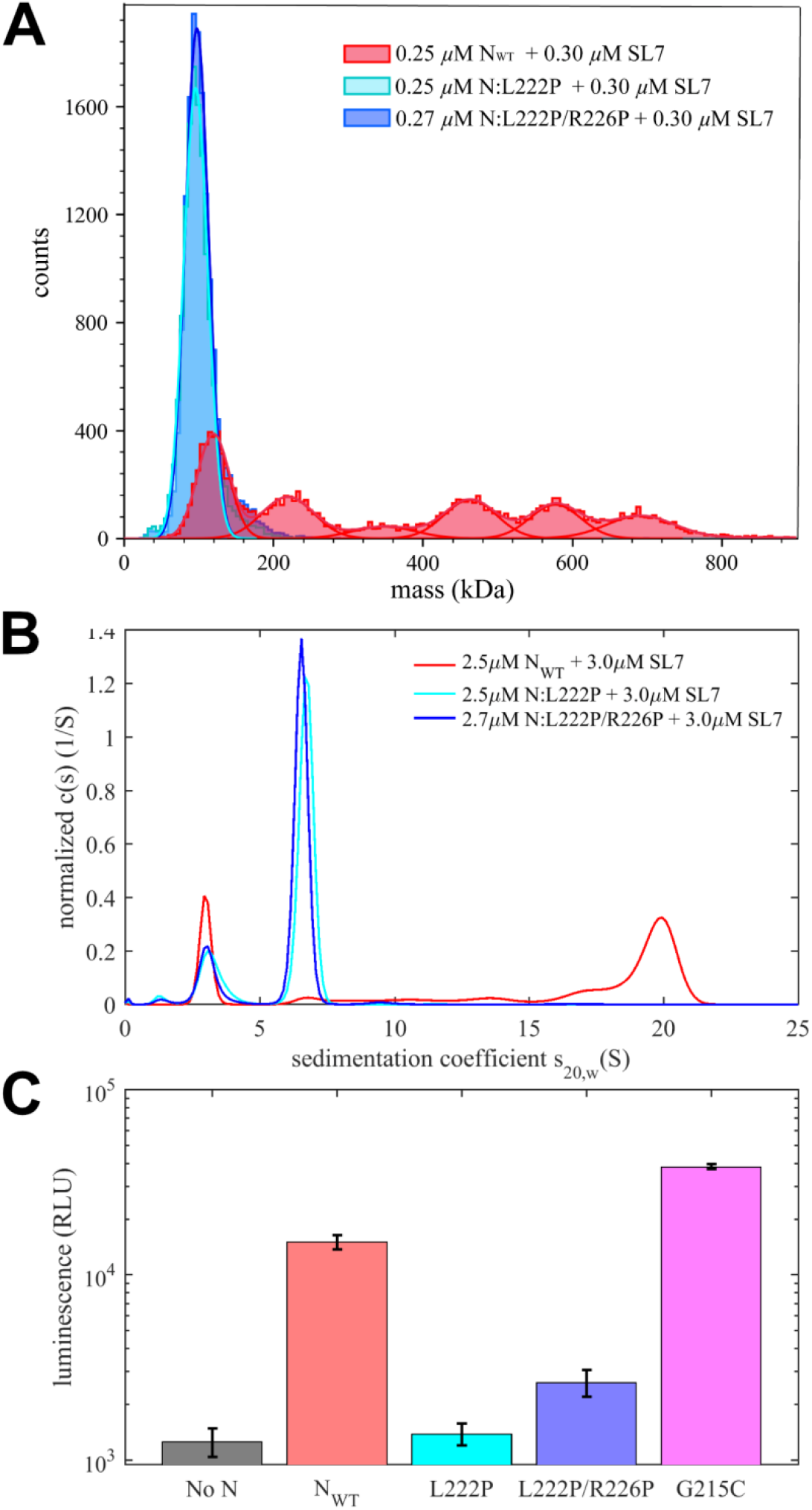
RNP formation of N-protein with SL7 depends on LRS oligomerization *in vitro* and in viral assembly. (A) Mass distributions from MP of mixtures of 0.25 µM N-protein with 0.3 µM SL7 in moderate ionic strength buffer B_65K_ for N_WT_ and the mutants inhibiting LRS oligomerization N:L222P and N:L222P/R226P. (B) Sedimentation coefficient distributions from SV-AUC experiments at 2.5 µM N-protein with 3.0 µM in buffer B_65K_ recorded at 260 nm. Shown are results with N_WT_, N:L222P and N:L222P/R226P. (C) Formation of VLPs containing SARS-CoV-2 structural proteins (S, E, M, and N) assembled in 293T cells with packaging RNA and a luciferase reporter. Efficiency of VLP formation was measured for different (or lacking) N-protein species indicated and quantified in relative luminescence units.

To study these mixtures at tenfold higher concentrations we turn to SV-AUC (Figure 5B), which for N_WT_ shows the familiar ≈20 S peak. By contrast, both LRS mutants only show drastically slower ≈6.5 S reaction boundaries. DLS data acquired for the same samples leads to hydrodynamic radii of 4.8 nm and 5.0 nm for the single and double mutant, respectively. The combination of sedimentation and diffusion coefficients yields molar mass estimates of 350 kDa for SL7 complexes with N:L222P, and 370 kDa for N:L222P/R226P. Thus, at low micromolar concentrations we can observe only complexes with 3 – 4 N-protein dimers bound to SL7, in contrast to the significantly larger and more compact 20 S complex with N_WT_ carrying intact LRS protein-protein interfaces.

It is useful to recapitulate the roles of the different assembly interfaces. The experiments contrasting LRS point mutants vs N_WT_ demonstrate the key role of LRS oligomerization in the assembly of large RNPs. However, clearly LRS oligomerization alone is not sufficient for the formation of large RNPs: When LRS oligomerization is switched on through binding of only short T_10_ NA ligands that are unable to scaffold N-protein, only ≈6 S complexes are formed at low micromolar concentrations, consistent with dimerization of dimers (Figure 2B, grey line) (17). This underlines the importance of simultaneous multivalent scaffolding for RNP formation. Even without LRS interactions, scaffolding of N-protein on NA can lead to intermediate size complexes at micromolar concentrations, as demonstrated in Figure 5B (blue and cyan lines) for binding to SL7 and in Figure 2 (blue and cyan lines) for binding to T_40_. However, these complexes are not very stable and dissociate at nanomolar concentrations, as shown in Figure 5A (blue and cyan histograms). We can conclude that both multivalent scaffolding and LRS oligomerization is necessary for large RNPs to form.

This can be observed clearly at nanomolar concentrations in MP experiments, where only joint interactions of NA-mediated scaffolding and LRS-mediated protein-protein oligomerization provide sufficient stability for the oligomers of {N dimer/2SL7} subunits to populate (Figure 5A, green). This can occur only if NA-mediated scaffolding acts to help cross-link {N dimer/2SL7} subunits, i.e., through shared interactions of different N dimers with the same SL7, or potentially through the same N-protein domain offering binding sites for multiple SL7 (as sketched in Figure 1H**/I**). Such additional {N dimer/2SL7} subunit interactions *via* the NA in concert with LRS interactions create the strong enhancement typical for avidity-based multi-valent interactions (in a simplistic picture through added binding energies). The same avidity-based enhancement of complex stability supports the formation of large RNP complexes at low micromolar concentrations, once scaffolding and LRS interactions act jointly.

### RNP complexes are stabilized through multivalent RNA binding of the CTD

We next focus on the question of what supports the formation of large RNPs with SL7 as NA substrate but not the similar sized T_40_. As shown in experiments with LRS mutants, both oligonucleotides allow scaffolding of multiple N-protein dimers, and in comparison with N_WT_ both have been shown to act in concert with LRS oligomerization. Nevertheless, RNP complexes with SL7 are roughly twice the size than those with T_40_. (A side-by-side comparison of T_40_ and SL7 complexes at the same concentration and under the experimental conditions of the RNP assay is provided in **Supplementary Figure S3**.)

Notwithstanding supporting roles of N-protein IDRs, the major nucleic acid binding sites are located in the folded NTD and CTD (14, 27, 28). Accordingly, we prepared constructs of isolated NTD and CTD domains to study their interactions with either T_40_ or SL7. By itself, NTD is monomeric and CTD dimeric due to its high-affinity dimerization interface; and neither shows detectable further self-association in the micromolar concentration range considered here, consistent with expectations from the literature (27, 29, 30). By applying low ionic strength conditions (B_10Na_) to drive the reaction into the maximal assembly state, we measured the size and composition of the CTD/NA and NTD/NA complexes by SV-AUC and obtain information on the protein footprint.

Figure 6A shows sedimentation coefficient distributions of 10 µM NTD in fivefold molar excess over T_40_ or SL7, respectively. For both oligonucleotides, a ≈5.2 S peak is observed with no or little free protein. The complex sediments much faster than either free NA or free NTD. Multi-signal analysis leads to an estimated protein/NA molar ratio of 5.4:1 on T_40_ and 4:1 – 5:1 on SL7. For T_40_, allowing for end effects, we can conclude a maximum for the footprint of ≈10 bases.

**Figure 6:**
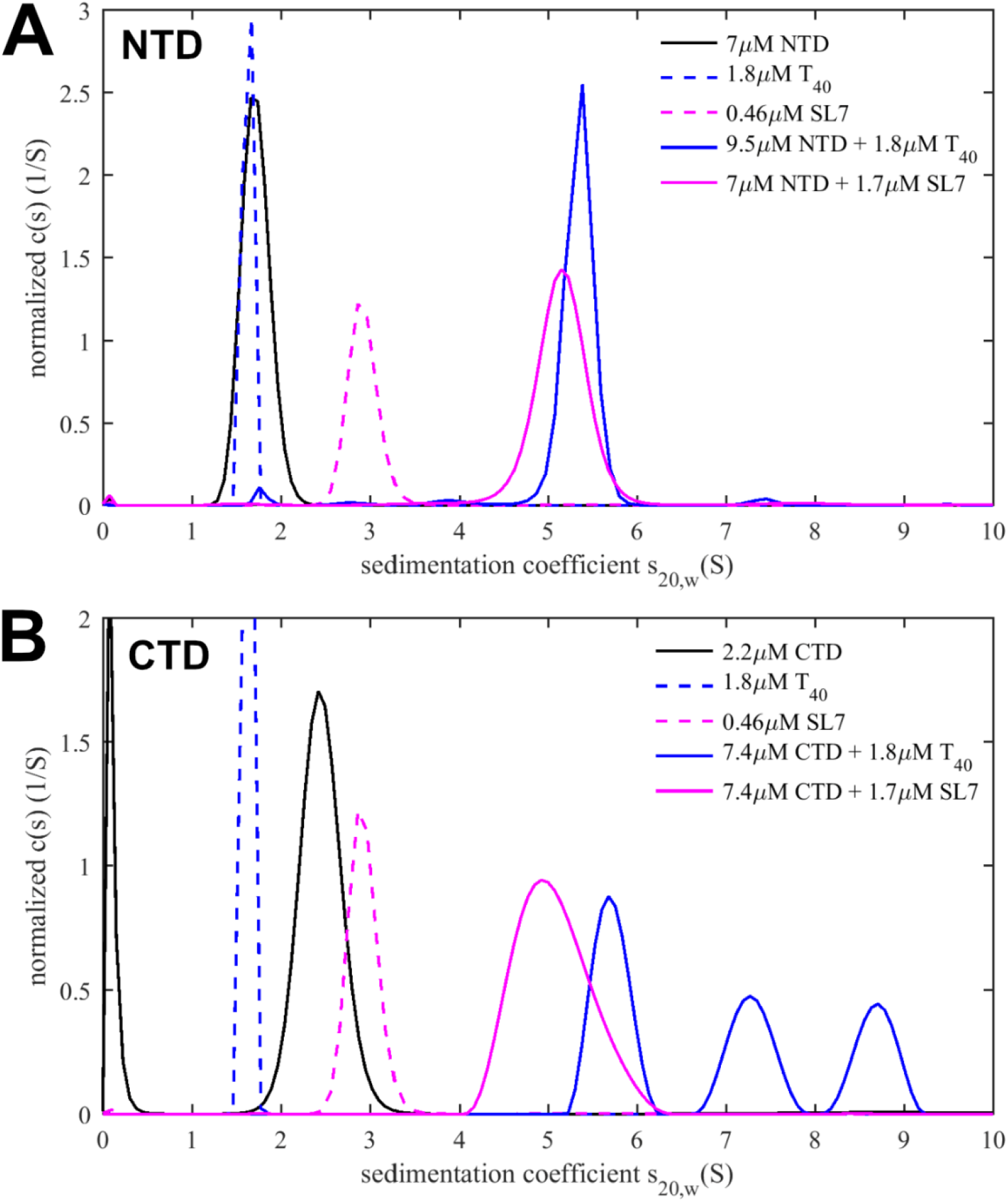
NTD and CTD binding to T_40_ and SL7. Sedimentation coefficient distributions *c*(*s*) from SV-AUC of NTD (A) or CTD (B) alone and in mixtures with T_40_ or SL7, respectively. Experiments are carried out in low ionic strength buffer B_10Na_ to promote formation of the complexes with maximum stoichiometry. Distributions of free T_40_ and SL7 are reduced by a factor 2. For NTD alone, the molecular weight determined from *c*(*s*) analysis and best-fit frictional ratio is 15.1 kDa, which compares well to the theoretically expected value of 15.2 kDa. For CTD alone, the experimental molecular weight is 27.3 kDa, which compares to the theoretical value of 26.6 kDa for a CTD dimer.

Analogous experiments for the CTD dimer are shown in Figure 6B. 10 µM CTD in fivefold molar excess over T_40_ in low ionic strength conditions produces similar sized CTD/T_40_ complexes compared to NTD/T_40_. By contrast, mixtures of CTD with SL7, under otherwise identical conditions, become cloudy upon mixing, and complexes of CTD/SL7 sediment significantly faster, up to ≈8.8 S. Based on the hydrodynamic scaling law s ∼ M^2/3^ one would expect the largest complex to have approximately twice the molar mass of the CTD/T_40_ complexes. Multi-signal analysis of all CTD/SL7 complexes results in an average molar ratio of ≈6 (±0.5):1 CTD per SL7, or 3 CTD dimers per SL7. This composition was used to determine the complex partial specific volume, and to determine *via* the Stokes equation the minimum mass of a hydrated spherical complex that can sediment as fast as 8.8 S, which is 140.8 kDa, or ≈181 kDa for more realistically assuming a moderately compact particle with frictional coefficient 1.3. This clearly exceeds the smallest possible complex with 6:1 stoichiometry (94.7 kDa for 3 CTD dimers + 1 SL7), but it would be consistent with a moderately asymmetric 12:2 complex (i.e., 6 CTD dimers binding 2 SL7).

Since SL7 does not oligomerize by itself, and the CTD does not oligomerize not beyond the dimer, we can conclude CTD dimers must exhibit at least a weak second site for SL7, to be able to accommodate two SL7 molecules as required for the large CTD/SL7 complexes observed. Mutual multi-valency of the CTD dimer and SL7 is consistent with the observed polydispersity of complexes as well as the aggregation propensity of these mixtures.

In conclusion, translated into the context of the FL N-protein, it appears the additional interactions of CTD dimer with SL7 on its second site act to weakly cross-link {N dimer/2SL7} and thereby stabilize the hexameric complexes of these subunits in cooperation with scaffolding and LRS protein-protein interactions (Figure 1H**/I**). When binding T_40_, these additional interactions are absent and only smaller complexes RNP complexes can form in the micromolar concentration range tested.

### The assembly of virus-like particles depends on the oligomerization in the LRS

To probe whether the involvement of LRS in the formation of RNPs *in vitro* extends to viral assembly *in vivo* we employed a recently described SARS-CoV-2 virus-like particle (VLP) assay (31). Briefly, the structural proteins S, E, M, and N are coexpressed in HEK 293T cells, and assembly of VLPs is initiated through co-transfection with a plasmid containing a 2 kb RNA packaging signal T20 along with a luciferase reporter transcript. VLPs from the supernatant of these producer cells are used to infect 293T receiver cells expressing SARS-CoV-2 entry factors ACE2 and TMPRSS2, and luminescence of the receiver cells is measured. Thus, introduction of mutant N-protein in this assay allows to test their impact on viral packaging and assembly efficiency (31).

Figure 5C shows a strong reduction of luminescence, relative to N_WT_, for the point mutants N:L222P and N:L222P/L226P, demonstrating that selectively inhibiting LRS interactions strongly impairs VLP formation. To test the opposite effect, we introduced the mutation N:G215C that was previously shown to enhance LRS interactions by stabilizing the helical state in the linker IDR (18). In the VLP assay, significantly elevated luminescence was observed for this enhancer mutation of LRS interactions (Figure 5C). This is consistent with the hypothesis that the protein-protein contacts from transient helices in the LRS and their coiled-coil oligomerization is an essential component of the assembly mechanism *in vivo*.

Since N-protein phosphorylation constitutes a switch between its intracellular and assembly functions (32), it is possible to increase the amount of assembly-competent N-protein in the VLP assay through abrogation of phosphorylation by treatment of the VLP producer cells with an inhibitor of GSK3 (CHIR98014), for which the SR-rich region of the linker provides abundant sites (5, 33–35). As shown previously, this results in significantly elevated VLP production (32). Introduction of the LRS mutants in the presence of this phosphorylation inhibitor led to a reduction of VLP assembly, but it reached a higher level than in the absence of the inhibitor (**Supplementary Figure S4**). Thus, it appears the structural destabilization of the LRS helices can be partially compensated in cells by increased intracellular levels of assembly-competent unphosphorylated N-protein.

## Discussion

The assembly pathway of SARS-CoV-2 N-protein to package and condense the large gRNA has remained enigmatic, despite cryo-ET density maps of distinct vRNP particles (1, 2), increasing atomistic insight in the structural and dynamic aspects of NA binding modes of the folded NTD and CTD domains (7, 14, 20, 30, 36, 37), and a recently described protein-protein interface in the disordered linker (17). In part this is due to the formidable problem of compounded effects of highly multivalent protein-NA and protein-protein interactions taking place side-by-side, the limited specificity of NA sites, high flexibility of N-protein imparted by the IDRs (38), and the resulting polymorphic nature of the complexes. Recent elegant work from the Morgan laboratory has suggested a model of six N-protein dimers organizing ≈600 bases of RNA in RNPs (16). In the present study, limited complex formation *in vitro* between model NAs and purified WT N-protein or point-mutants, along with complementary VLP assembly experiments, revealed further architectural principles of RNPs that contribute to a more complete picture of assembly.

It is likely that simultaneous binding of N-protein dimers to the 5’ untranslated region of the gRNA, *via* NTD likely preferring unpaired single stranded regions and CTD preferring double stranded RNA, creates locally a high density of N-protein in close proximity (7, 9, 11, 39). As the NA-bound N-protein NTD induces a conformational change in the LRS to a self-assembly-competent helical state (17), we propose the cooperative oligomerization of LRS helices “rolls up” N-protein dimer complexes into hexameric assemblies with LRS oligomers on the inside, similar to the sketch Figure 1I. There appears to be a role for macromolecular condensates to help initiate and/or propagate vRNP formation by ensuring high local N-protein concentrations (9, 11–13); and plausibly for the same ultra-weak attractive interactions that can lead to macromolecular condensates to help holding semi-liquid high-density states together (16).

Our *in vitro* data on the assembly properties of LRS point mutant N-proteins clearly demonstrate LRS assembly is a key mechanism, without which only small complexes can form even at high concentrations resulting from simple scaffolding of N-protein on the NA. The protein-protein interface in the LRS allows such NA-bound N-protein dimers to further self-assemble into three-dimensional structures. This is consistent with optical tweezer experiments by Morse et al. examining multiple copies of N-protein binding to a single molecule of ssDNA, which show a cooperative reorganization of an initially formed protein/DNA complex (40) that we would attribute to LRS oligomerization after N-protein scaffolding on NA.

In support that LRS oligomerization is essential also for assembly *in vivo*, we carried out VLP experiments that demonstrate strongly reduced or enhanced VLP formation as a result of N-protein point mutants that diminish or enhance coiled-coil oligomerization of LRS helices, respectively. Even though the packaged NA in the VLPs is different from the longer viral genome, and different from the stem-loop based *in vitro* model system for RNP assembly, LRS assumes a critical role in both. Several lines of evidence suggest this conclusion also extends to live SARS-CoV-2 virus: While we have shown previously that LRS oligomerization occurs similarly in N-protein of related coronaviruses, SARS-CoV-2 Delta variant clades containing the G215C mutation that enhances LRS oligomerization dramatically outperformed those without G215C (18). Conversely, mutations of the LRS that destabilize the oligomerization of LRS helices are excluded from the exhaustively sampled mutational landscape of viable SARS-CoV-2 (17).

Interestingly, the formation of large RNPs in the *in vitro* model examined here depends, besides the LRS, also on multivalent binding of RNA by the CTD. In experiments focused on the binding of the isolated CTD domain to either T_40_ or SL7, the major difference was the assembly of significantly larger complexes with SL7, which are of a size and composition that require the ability of a CTD dimer to bind two SL7 molecules. While a second binding site for RNA on the CTD dimer is unexpected, additional support for at least a weak second RNA site on the CTD dimer comes from our observed turbidity upon mixing of CTD with SL7 (which depends on mutually multivalent interactions for either aggregation or LLPS), similarly observed aggregation at high concentration in the presence of stem-loop S2m (28). It would also be consistent with multi-step binding of isolated CTD to DNA observed in optical tweezer experiments by Morse *et al.* (40), and conclusions from structural modeling by Padroni *et al.* (36). In the present RNP assembly assay with FL-N, this appears to be the key difference between the different NA substrates, where, similar to the CTD results, binding of T_40_ leads to only moderately sized complexes, whereas SL7 supports significantly larger RNP assemblies. Mechanistically, since the second SL7 binding site on the CTD dimer acts in concert with LRS oligomerization, the resulting avidity can amplify even a weak second SL7 site to provide significant complex stabilization. We speculate that in the interaction with gRNA the ability to accommodate a second RNA interaction at the CTD dimer may similarly stabilize compaction and scaffolding of stem loops on N-protein dimers which would then enhance LRS oligomerization and vRNP formation.

An important aspect of the assembly model is the existence of an upper limit of size of the RNP complexes to form discrete particles. Combining SV-AUC and DLS at a range of micromolar concentrations, we found this limit to be at a hexamer of N-protein dimers, consistent with results from Carlson *et al.* by MP in dilute solution after chemical crosslinking (16). From cryo-ET data, vRNPs are heterogeneous, but a subclass appear to be hexagonally packed (2). The previously observed pleomorphic ability of LRS helices to form different oligomeric states (17) may contribute to the heterogeneity of particles. Unfortunately it is difficult to reference assembly models in detail to the electron microscopy density maps due to the large fractions of IDRs of N-protein that are highly flexible. For example, end-to-end distances of the linker IDR alone may range from 2 to 12 nm (38). The resulting variation of the local conformation may cause lack of contrast in maps averaged across vRNP particles. Besides the configuration of the N-protein scaffold, it is also unclear in which way gRNA is packaged onto the vRNPs and contributes to its structure. Various secondary structural features involve the majority of gRNA (41), and N-protein flexibility may contribute structural plasticity to accommodate different RNA structural elements. On the other hand, ssRNA is also highly flexible with a persistence length of ≈2 nm (42), and RNA 3D structures may not be unique (43), so that it is also conceivable that RNA structures may be dynamically reshaped by N-protein interactions (44, 45).

At least a dimensional analysis is of interest. As pointed out by Carlson *et al.*, if the size of the genome is divided by the number of vRNPs, on average approximately 800 nt would be scaffolded by each vRNP (16), or somewhat less accounting for stretches connecting neighboring particles. This could be achieved if a hexamer of N-protein dimers would on average bind 60-65 nt per N-protein, which is not dissimilar to the RNP complexes obtained in the SL7 stem-loop model system in the present work, or the SL8 and other stem-loop model examined by Carlson *et al.* (16). Thus assembly of N-protein dimer/RNA stem-loop configurations as depicted in Figure 1H**/I** could theoretically roughly satisfy scaffolding requirements of vRNPs. Furthermore, for the RNP particles with SL7 we measured a Stokes radius of 8.5 nm, which seems reasonably consistent with the 14-16 nm sized structures observed in cryo-ET (1, 2).

The pleomorphic ability of vRNPs may extent to its architectural principles. Adly *et al.* and Syed *et al.* have recently shown by MP, EM, and in the VLP assay that truncated N-protein N:Δ(1–209) can still form RNPs similar in size and shape to FL-N (32, 46). Remarkably, this is despite lacking (in addition to the N-arm IDR and the SR-rich region of the linker IDR) the NTD, which is thought to confer specificity of the assembly (7, 39). However, the LRS region was critical for assembly of N:Δ(1–209) (46), consistent with our model of LRS oligomerization as a central mechanism for vRNP formation. N:Δ(1–209) arises as an alternate viral transcript due to the introduction of a new transcription regulatory sequence associated with the R203K/G204R mutation in Alpha, Gamma, and Omicron variants. Interestingly, a significant increase in the number of RNPs was found by cryo-ET in Omicron variants (47), suggesting altered RNA packaging. However, other factors may also contribute to this, such as new protein-protein interactions between Omicron N-arm IDRs arising from the obligatory mutation P13L of Omicron variants (48) and possibly epistatic interactions with other N-protein mutations that coexist in variants of concern. In light of the pleomorphic architectures of vRNP, it appears that the interfaces in LRS oligomerization along with the NA binding sites on the CTD would remain as the most promising antiviral targets.

Finally, from a methodological point of view, the current work highlights the strong complementarity between the relatively new method of MP and the classical biophysical methods of SV-AUC and DLS for studying interacting systems, all of which report on distributions of macromolecules and their complexes in equilibrium in solution without requiring separations, modifications, or surface immobilization. Their combination provides a powerful new approach for the study of multi-step assembly processes. MP has superior mass resolution and applications to the analysis of interacting systems have been pioneered (49), but the technique is limited to nanomolar concentrations and species above 30 kDa. In the present case it served well to study onset of the multi-step assembly reaction. SV-AUC far extends the concentration range and therefore allowed observation of the assembly up to the largest assembly products, adding spectral resolution to distinguish composition of the protein/NA complexes. (Fluorescence detected SV-AUC also extends sensitivity into the picomolar range, but attachment of a fluorescent tag is be detrimental in the present system (19).) Importantly, in the intermediate concentration range accessible to both MP and SV-AUC we obtained highly consistent results. While SV-AUC carries molar mass information only indirectly, folded into hydrodynamic migration, the latter can be decomposed into size and mass information with the help of independent information on translational friction, here taken from DLS for the largest reaction product. Alternatively, shape information may be derived by combination of the complex mass values from MP with sedimentation coefficients from SV. Furthermore, an additional promising aspect from this combination of methods is that the notorious dependency of sedimentation boundary patterns on the complex lifetimes in SV-AUC may be untangled with information from MP. Jointly, MP, SV-AUC, and DLS should be amenable to global multi-method analysis (50) but detailed affinities of all assembly steps would be of limited interest in the current case due to the pleomorphic nature of the RNP assembly.

## Acknowledgements

We thank Vanessa Wall, Brianna Higgins, and J-P Denson for N-CTD protein production. This project was funded in part with federal funds from the National Cancer Institute, National Institutes of Health Contract 75N91019D00024. J.A.D got support from by a grant from the NIH (R21AI59666) and by support from the Howard Hughes Medical Institute and the Gladstone Institutes. M.O. received support from NIH Host pathogen mapping initiative (HPMI) grant (U19 AI135990), the James B. Pendleton Charitable Trust, the Roddenberry Foundation, P. and E. Taft, and the Gladstone Institutes. M.O. is a Chan Zuckerberg Biohub – San Francisco Investigator. This work was supported by the Intramural Research Programs of NIBIB, NHLBI, NIDDK, and NCI of the National Institutes of Health, Bethesda.

## Materials and Methods

### Protein expression and purification

Wildtype and mutant full-length SARS-CoV-2 N-proteins were expressed and purified as described previously (17, 18). Briefly, the full-length protein was cloned into the pET-29a(+) expression vector fused to DNA encoding a 6xHis tag followed by Tobacco Etch Virus (TEV) cleavage site at N-terminus, and transformed into One Shot BL21(DE3)pLysS E. coli (Thermo Fisher Scientific, Carlsbad, CA). After cell lysis, the protein was bound to by Ni-NTA column, followed by purification with unfolded and refolded steps to remove residual protein-bound bacterial nucleic acid (11). After Ni^2+^ affinity chromatography, the eluate was subject to cleavage of the 6xHis tag by TEV and purification by affinity and size exclusion chromatography. 95% purity of the proteins was confirmed by SDS-PAGE. The ratio of absorbance at 260 nm and 280 nm of ∼0.50-0.55 confirmed absence of nucleic acid.

SARS-CoV-2 N-protein NTD (48–173) was cloned into kanamycin-resistant, NdeI/XhoI-digested plasmid pET29a vector (GenScript) with 6xHis tag included at the N-terminus. After expression and lysis, NTD was purified by Ni-NTA chromatography. The C-terminal domain (247–364) was generated as previously described (51): Briefly, an N-terminal 6xHis tagged N-CTD construct preceded by a TEV protease cleavage site was expressed in E. cli BL21(DE3) using Dynamite Broth as described in (52). After expression and lysis, protein was purified by Ni^2+^ affinity chromatography followed by cleavage of the tag and purification by affinity and size exclusion chromatography. Protein purity was validated by SDS-PAGE chromatography and electrospray ionization mass spectrometry The oligonucleotide T_40_ and stem-loop RNA SL7 were purchased from Integrated DNA Technologies (Skokie, IL), as purified by HPLC and lyophilized. After reconstitution, to obtain optimal RNA secondary structure, SL7 was subject to thermal denaturation at 95 °C for 2 min followed by slowly cooling to room temperature over 1-2 hrs.

Prior to experiments, protein and nucleic acid samples were dialyzed into working buffer and concentrations were determined by UV-Vis spectrophotometry. More detailed information on all protein constructs can be found in the **Supplementary Materials and Methods**.

### Virus-like particle assay

A virus-like particle (VLP) assay was employed as a physiological model to test the efficiency of packaging and assembly as a function of the mutations on SARS-CoV-2 N protein (31). Implementation in the present work mirrors the detailed description in (46). Briefly, the VLPs were generated by co-expressing all four structural proteins of SARS-CoV-2 in HEK 293T cells. A 2 kb viral packaging signal was incorporated into the untranslated region of a luciferase reporter plasmid, resulting in the transcripts to be packaged in the VLPs, which were added to receiver 293T cells expressing ACE2 and TMPRSS2.

Luminescence in receiver cells was measured using a Promega Luciferase Assay System (Promega E1501). More details can be found in the **Supplementary Materials and Methods**.

### Mass Photometry

Mass photometry experiments were carried out using a TwoMP instrument (Refeyn, UK). Samples were loaded in the mini-wells created by a silicone gasket placed on top of a microscope coverslip on the microscope stage. First, the samples were prepared by mixing the stock solutions with the working buffers in the Eppendorf tubes. Prior to MP data acquisition, the same working buffer (10-15 μL) was loaded into the mini-well to focus the objective, after which 1 or 2 μL of the sample was added to the buffer droplet, mixed and measured immediately.

The TwoMP instrument was calibrated with Beta-Amylase from Sweet Potato (Sigma A8781) and Thyroglobulin from Bovine Thyroid (Sigma T9145) as recommended by the manufacturer. Mass photometry data was acquired with AcquireMP software and the analysis was performed with DiscoverMP software (Refeyn, UK). For each MP data set, histograms of individual mass measurements were inspected, and the mass distribution was fitted with Gaussian curves to estimate the average molar mass of the selected distributions.

### Sedimentation Velocity Analytical Ultracentrifugation

Sedimentation velocity analytical ultracentrifugation (SV-AUC) experiments were performed in a ProteomeLab Xl-I analytical ultracentrifuge (Beckman Coulter, Indianapolis, IN) by following the standard protocol (53). The macromolecules and their mixtures were loaded in 12- or 3-mm charcoal-filled Epon double-sector centerpieces with sapphire windows. The AUC cell assemblies filled with samples were inserted into An-50 or An-60 rotors, followed by temperature equilibration at 20 °C in the AUC. Data were acquired with Rayleigh interference optics and absorbance optics at 260 nm and/or 280 nm, depending on the solution composition. Instrument calibration factors were determined as previously described (54). The SV-AUC data were analyzed using sedimentation coefficient distribution *c*(*s*) model in the software SEDFIT (55) (https://sedfitsedphat.nibib.nih.gov/software).

Multi-signal analysis was based on the analysis of integrated *c*(*s*) peaks obtained in analyses of families of sedimentation profiles recorded with different absorbance and refractometric optical signals. Signal increments were determined from the signals in the sedimentation boundaries of free NA or protein species, respectively, quantified through the integration of the respective *c*(*s*) distribution peaks and their known concentration. Partial-specific volumes of complexes were calculated as weight-averages, based in a partial-specific volume of RNA of 0.59 mL/g in potassium salts (56). Partial-specific volumes of protein species, and calculations using Stokes and Svedberg equations were carried out using the calculator functions in SEDFIT.

### Dynamic Light Scattering

Autocorrelation functions (ACFs) of the samples were collected in either a NanoStar instrument (Wyatt Technology, Santa Barbara, CA) or Prometheus Panta (Nanotemper, Germany) instrument at 20°C. In NanoStar, 20 µL samples were inserted into a 1 µL quartz cuvette (WNQC01-00, Wyatt Instruments).

Laser light scattering was measured at 658 nm at detection angle of 90°. In Prometheus Panta, the samples were loaded into a capillary (Nanotemper PR-AC002) and ACFs were acquired using the 405 nm laser at the detection angle of 140°. The ACFs from both instruments were processed and analyzed in SEDFIT.

## References

1. Klein, S., Cortese, M., Winter, S.L., Wachsmuth-Melm, M., Neufeldt, C.J., Cerikan, B., Stanifer, M.L., Boulant, S., Bartenschlager, R. and Chlanda, P. (2020) SARS-CoV-2 structure and replication characterized by in situ cryo-electron tomography. Nat. Commun., 11, 5885.

2. Yao, H., Song, Y., Chen, Y., Wu, N., Xu, J., Sun, C., Zhang, J., Weng, T., Zhang, Z., Wu, Z., et al. (2020) Molecular Architecture of the SARS-CoV-2 Virus. Cell, 183, 730–738.e13.

3. Wu, W., Cheng, Y., Zhou, H., Sun, C. and Zhang, S. (2023) The SARS-CoV-2 nucleocapsid protein: its role in the viral life cycle, structure and functions, and use as a potential target in the development of vaccines and diagnostics. Virol. J., 20, 6.

4. Yu, H., Guan, F., Miller, H., Lei, J. and Liu, C. (2023) The role of SARS-CoV-2 nucleocapsid protein in antiviral immunity and vaccine development. Emerg. Microbes Infect., 12.

5. Schuck, P. and Zhao, H. (2023) Diversity of Short Linear Interaction Motifs in SARS-CoV-2 Nucleocapsid Protein. MBio, in press.

6. Masters, P.S. (2019) Coronavirus genomic RNA packaging. Virology, 537, 198–207.

7. Korn, S.M., Dhamotharan, K., Jeffries, C.M. and Schlundt, A. (2023) The preference signature of the SARS-CoV-2 Nucleocapsid NTD for its 5’-genomic RNA elements. Nat. Commun., 14, 3331.

8. Savastano, A., Ibáñez de Opakua, A., Rankovic, M. and Zweckstetter, M. (2020) Nucleocapsid protein of SARS-CoV-2 phase separates into RNA-rich polymerase-containing condensates. Nat. Commun., 11, 6041.

9. Roden, C.A., Dai, Y., Giannetti, C.A., Seim, I., Lee, M., Sealfon, R., McLaughlin, G.A., Boerneke, M.A., Iserman, C., Wey, S.A., et al. (2022) Double-stranded RNA drives SARS-CoV-2 nucleocapsid protein to undergo phase separation at specific temperatures. Nucleic Acids Res., 50, 8168–8192.

10. Cascarina, S.M. and Ross, E.D. (2022) Phase Separation by the SARS-CoV-2 Nucleocapsid Protein: Consensus and Open Questions. J. Biol. Chem., 10.1016/j.jbc.2022.101677.

11. Carlson, C.R., Asfaha, J.B., Ghent, C.M., Howard, C.J., Hartooni, N., Safari, M., Frankel, A.D. and Morgan, D.O. (2020) Phosphoregulation of Phase Separation by the SARS-CoV-2 N Protein Suggests a Biophysical Basis for its Dual Functions. Mol. Cell, 80, 1092–1103.e4.

12. Cubuk, J., Alston, J.J., Incicco, J.J., Singh, S., Stuchell-Brereton, M.D., Ward, M.D., Zimmerman, M.I., Vithani, N., Griffith, D., Wagoner, J.A., et al. (2021) The SARS-CoV-2 nucleocapsid protein is dynamic, disordered, and phase separates with RNA. Nat. Commun., 12, 1936.

13. Iserman, C., Roden, C.A., Boerneke, M.A., Sealfon, R.S.G., McLaughlin, G.A., Jungreis, I., Fritch, E.J., Hou, Y.J., Ekena, J., Weidmann, C.A., et al. (2020) Genomic RNA Elements Drive Phase Separation of the SARS-CoV-2 Nucleocapsid. Mol. Cell, 80, 1078–1091.

14. Pontoriero, L., Schiavina, M., Korn, S.M., Schlundt, A., Pierattelli, R. and Felli, I.C. (2022) NMR Reveals Specific Tracts within the Intrinsically Disordered Regions of the SARS-CoV-2 Nucleocapsid Protein Involved in RNA Encountering. Biomolecules, 12, 929.

15. Cubuk, J., Alston, J.J., Incicco, J.J., Holehouse, A.S., Hall, K.B., Stuchell-Brereton, M.D. and Soranno, A. (2023) The disordered N-terminal tail of SARS CoV-2 Nucleocapsid protein forms a dynamic complex with RNA. bioRxiv, 10.1101/2023.02.10.527914.

16. Carlson, C.R., Adly, A.N., Bi, M., Howard, C.J., Frost, A., Cheng, Y. and Morgan, D.O. (2022) Reconstitution of the SARS-CoV-2 ribonucleosome provides insights into genomic RNA packaging and regulation by phosphorylation. J. Biol. Chem., 298, 102560.

17. Zhao, H., Wu, D., Hassan, S.A., Nguyen, A., Chen, J., Piszczek, G. and Schuck, P. (2023) A conserved oligomerization domain in the disordered linker of coronavirus nucleocapsid proteins. Sci. Adv., 9, eadg6473.

18. Zhao, H., Nguyen, A., Wu, D., Li, Y., Hassan, S.A., Chen, J., Shroff, H., Piszczek, G. and Schuck, P. (2022) Plasticity in structure and assembly of SARS-CoV-2 nucleocapsid protein. PNAS Nexus, 1, pgac049.

19. Zhao, H., Wu, D., Nguyen, A., Li, Y., Adão, R.C., Valkov, E., Patterson, G.H., Piszczek, G. and Schuck, P. (2021) Energetic and structural features of SARS-CoV-2 N-protein co-assemblies with nucleic acids. iScience, 24, 102523.

20. Dinesh, D.C., Chalupska, D., Silhan, J., Koutna, E., Nencka, R., Veverka, V. and Boura, E. (2020) Structural basis of RNA recognition by the SARS-CoV-2 nucleocapsid phosphoprotein. PLOS Pathog., 16, e1009100.

21. Schuck, P. and Zhao, H. (2017) Sedimentation Velocity Analytical Ultracentrifugation: Interacting Systems CRC Press, Boca Raton, FL.

22. Balbo, A., Minor, K.H., Velikovsky, C.A., Mariuzza, R.A., Peterson, C.B. and Schuck, P. (2005) Studying multiprotein complexes by multi-signal sedimentation velocity analytical ultracentrifugation. Proc. Natl. Acad. Sci. USA, 102, 81–86.

23. Schuck, P. (2010) Sedimentation patterns of rapidly reversible protein interactions. Biophys. J., 98, 2005–2013.

24. Record, M.T., Anderson, C.F. and Lohman, T.M. (1978) Thermodynamic analysis of ion effects on the binding and conformational equilibria of proteins and nucleic acids: the roles of ion association or release, screening, and ion effects on water activity. Q. Rev. Biophys., 11, 103–178.

25. Li, Y., Struwe, W.B. and Kukura, P. (2020) Single molecule mass photometry of nucleic acids. Nucleic Acids Res., 48, E97.

26. Casassa, E.F. and Eisenberg, H. (1961) Partial specific volumes and refractive index increments in multicomponent systems. J. Phys. Chem., 65, 427–433.

27. Chang, C.K., Hou, M.H., Chang, C.F., Hsiao, C.D. and Huang, T.H. (2014) The SARS coronavirus nucleocapsid protein - Forms and functions. Antiviral Res., 103, 39–50.

28. Dang, M. and Song, J. (2022) CTD of SARS-CoV-2 N protein is a cryptic domain for binding ATP and nucleic acid that interplay in modulating phase separation. Protein Sci., 31, 345–356.

29. Wu, C., Qavi, A.J., Hachim, A., Kavian, N., Cole, A.R., Moyle, A.B., Wagner, N.D., Sweeney-Gibbons, J., Rohrs, H.W., Gross, M.L., et al. (2021) Characterization of SARS-CoV-2 nucleocapsid protein reveals multiple functional consequences of the C-terminal domain. iScience, 24, 102681.

30. Zinzula, L., Basquin, J., Bohn, S., Beck, F., Klumpe, S., Pfeifer, G., Nagy, I., Bracher, A., Hartl, F.U. and Baumeister, W. (2021) High-resolution structure and biophysical characterization of the nucleocapsid phosphoprotein dimerization domain from the Covid-19 severe acute respiratory syndrome coronavirus 2. Biochem. Biophys. Res. Commun., 538, 54–62.

31. Syed, A.M., Taha, T.Y., Tabata, T., Chen, I.P., Ciling, A., Khalid, M.M., Sreekumar, B., Chen, P.-Y., Hayashi, J.M., Soczek, K.M., et al. (2021) Rapid assessment of SARS-CoV-2-evolved variants using virus-like particles. Science, 374, 1626–1632.

32. Syed, A.M., Ciling, A., Chen, I.P., Carlson, C.R., Adly, A., Martin, H., Taha, T.Y., Khalid, M.M., Bouhaddou, M., Ummadi, M., et al. (2023) SARS-CoV-2 evolution balances conflicting roles of N protein phosphorylation. Available SSRN https://ssrn.com/abstract=4472729, 10.2139/ssrn.4472729.

33. Johnson, B.A., Zhou, Y., Lokugamage, K.G., Vu, M.N., Bopp, N., Crocquet-Valdes, P.A., Kalveram, B., Schindewolf, C., Liu, Y., Scharton, D., et al. (2022) Nucleocapsid mutations in SARS-CoV-2 augment replication and pathogenesis. PLOS Pathog., 18, e1010627.

34. Yaron, T.M., Heaton, B.E., Levy, T.M., Johnson, J.L., Jordan, T.X., Cohen, B.M., Kerelsky, A., Lin, T., Liberatore, K.M., Bulaon, D.K., et al. (2022) Host protein kinases required for SARS-CoV-2 nucleocapsid phosphorylation and viral replication. Sci. Signal., 15, 1–17.

35. Liu, X., Verma, A., Garcia, G., Ramage, H., Lucas, A., Myers, R.L., Michaelson, J.J., Coryell, W., Kumar, A., Charney, A.W., et al. (2021) Targeting the coronavirus nucleocapsid protein through GSK-3 inhibition. Proc. Natl. Acad. Sci. U. S. A., 118, 1–9.

36. Padroni, G., Bikaki, M., Novakovic, M., Wolter, A.C., Rüdisser, S.H., Gossert, A.D., Leitner, A. and Allain, F.H.T. (2023) A hybrid structure determination approach to investigate the druggability of the nucleocapsid protein of SARS-CoV-2. Nucleic Acids Res., 51, 4555–4571.

37. Redzic, J.S., Lee, E., Born, A., Issaian, A., Henen, M.A., Nichols, P.J., Blue, A., Hansen, K.C., D’Alessandro, A., Vögeli, B., et al. (2021) The Inherent Dynamics and Interaction Sites of the SARS-CoV-2 Nucleocapsid N-Terminal Region. J. Mol. Biol., 433, 167108.

38. Ró?ycki, B. and Boura, E. (2022) Conformational ensemble of the full-length SARS-CoV-2 nucleocapsid (N) protein based on molecular simulations and SAXS data. Biophys. Chem., 288, 106843.

39. Estelle, A.B., Forsythe, H.M., Yu, Z., Hughes, K., Lasher, B., Allen, P., Reardon, P.N., Hendrix, D.A. and Barbar, E.J. (2023) RNA structure and multiple weak interactions balance the interplay between RNA binding and phase separation of SARS-CoV-2 nucleocapsid. PNAS Nexus, 2, 1–31.

40. Morse, M., Sefcikova, J., Rouzina, I., Beuning, P.J. and Williams, M.C. (2022) Structural domains of SARS-CoV-2 nucleocapsid protein coordinate to compact long nucleic acid substrates. Nucleic Acids Res., 10.1093/nar/gkac1179.

41. Cao, C., Cai, Z., Xiao, X., Rao, J., Chen, J., Hu, N., Yang, M., Xing, X., Wang, Y., Li, M., et al. (2021) The architecture of the SARS-CoV-2 RNA genome inside virion. Nat. Commun., 12, 1–14.

42. Chen, H., Meisburger, S.P., Pabit, S.A., Sutton, J.L., Webb, W.W. and Pollack, L. (2012) Ionic strength- dependent persistence lengths of single-stranded RNA and DNA. Proc. Natl. Acad. Sci. U. S. A., 109, 799–804.

43. Ding, J., Lee, Y.T., Bhandari, Y., Schwieters, C.D., Fan, L., Yu, P., Tarosov, S.G., Stagno, J.R., Ma, B., Nussinov, R., et al. (2023) Visualizing RNA conformational and architectural heterogeneity in solution. Nat. Commun., 14.

44. Perlmutter, J.D. and Hagan, M.F. (2015) Mechanisms of virus assembly. Annu. Rev. Phys. Chem., 66, 217–239.

45. Larson, S.B. and McPherson, A. (2001) Satellite tobacco mosaic virus RNA: Structure and implications for assembly. Curr. Opin. Struct. Biol., 11, 59–65.

46. Adly, A.N., Bi, M., Carlson, C.R., Syed, A.M., Ciling, A., Doudna, J.A., Cheng, Y. and Morgan, D.O. (2023) Assembly of SARS-CoV-2 ribonucleosomes by truncated N* variant of the nucleocapsid protein. J. Biol. Chem., in press, 105362.

47. Ma, X., Wang, Y., Gao, Y., Wang, Y., Yan, A., Chen, J., Zhang, L., Wang, P., Zhao, J. and Liu, Z. (2023) Structural Characteristics of the SARS-CoV-2 Omicron lineages BA.1 and BA.2 virions. Signal Transduct. Target. Ther., 8, 131.

48. Nguyen, A., Zhao, H., Myagmarsuren, D., Srinivasan, S., Wu, D., Chen, J., Piszczek, G. and Schuck, P. (2023) Modulation of Biophysical Properties of Nucleocapsid Protein in the Mutant Spectrum of SARS-CoV-2. bioRxiv, 10.1101/2023.11.21.568093.

49. Wu, D. and Piszczek, G. (2020) Measuring the affinity of protein-protein interactions on a single-molecule level by mass photometry. Anal. Biochem., 592, 113575.

50. Zhao, H. and Schuck, P. (2012) Global multi-method analysis of affinities and cooperativity in complex systems of macromolecular interactions. Anal. Chem., 84, 9513–9519.

51. Karkanitsa, M., Li, Y., Valenti, S., Spathies, J., Kelly, S., Mehalko, J., Drew, M., Denson, J., Putman, Z. and Fathi, P. (2023) Dynamics of SARS-CoV-2 seroprevalence in a large US population over a period of 12 months. medRxiv, 10.1101/2023.10.20.23297329.

52. Taylor, T., Denson, J.-P. and Esposito, D. (2017) Optimizing Expression and Solubility of Proteins in E. coli Using Modified Media and Induction Parameters. Methods Mol. Biol., 1586, 65–82.

53. Schuck, P., Zhao, H., Brautigam, C.A. and Ghirlando, R. (2015) Basic Principles of Analytical Ultracentrifugation CRC Press, Boca Raton, FL.

54. Ghirlando, R., Balbo, A., Piszczek, G., Brown, P.H., Lewis, M.S., Brautigam, C.A., Schuck, P. and Zhao, H. (2013) Improving the thermal, radial, and temporal accuracy of the analytical ultracentrifuge through external references. Anal. Biochem., 440, 81–95.

55. Schuck, P. (2016) Sedimentation Velocity Analytical Ultracentrifugation: Discrete Species and Size-Distributions of Macromolecules and Particles CRC Press, Boca Raton, FL.

56. Greive, S.J., Lins, A.F. and von Hippel, P.H. (2005) Assembly of an RNA-protein complex. Binding of NusB and NusE (S10) proteins to boxA RNA nucleates the formation of the antitermination complex involved in controlling rRNA transcription in Escherichia coli. J. Biol. Chem., 280, 36397-36408.

