## Supplementary Figure S2 for "Assembly reactions of SARS-CoV-2 nucleocapsid protein with nucleic acid"

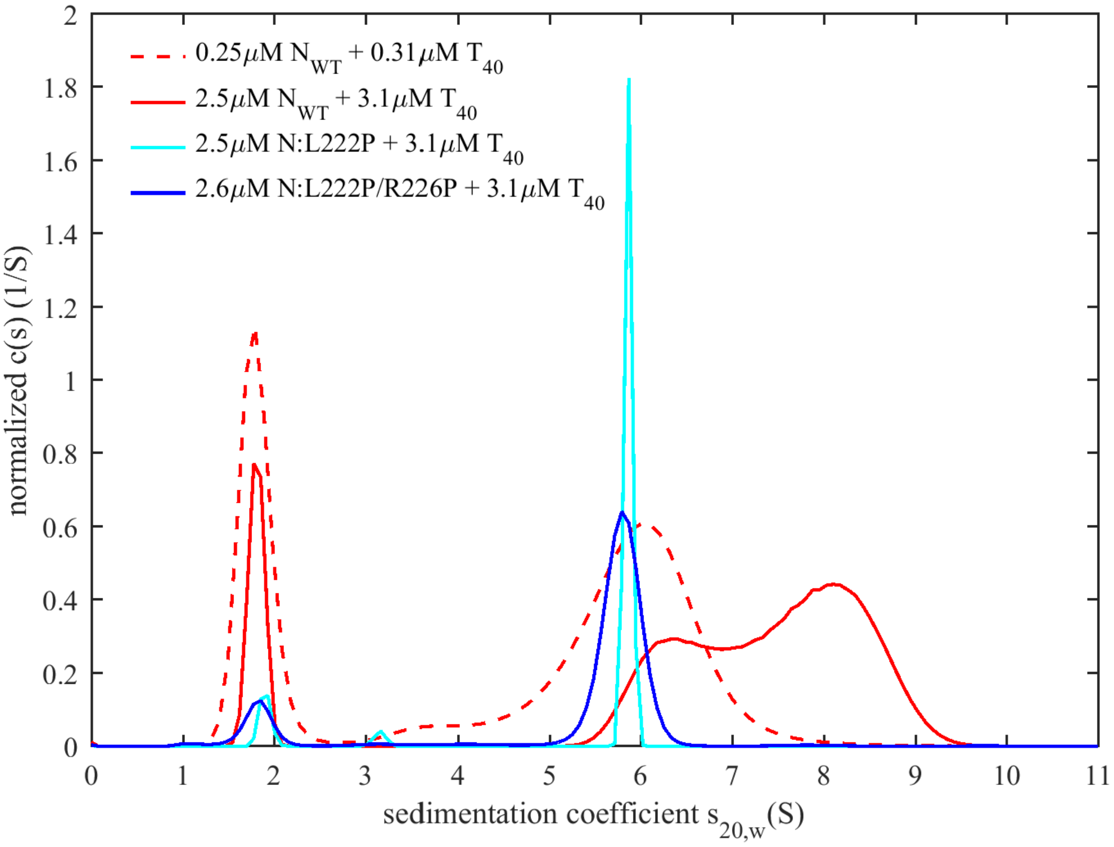


**Supplementary Figure S2: Simultaneous LRS oligomerization and scaffolding on the oligonucleotide $T_{40}$ in moderate ionic strength buffer $B_{65K}$.** Shown are sedimentation coefficient distributions recorded at 260 nm for $N_{WT}$ (red) and the LRS mutants N:L222P (cyan, magnitude reduced by factor 5) and N:L222P/R226P (blue, magnitude reduced by factor 2) at concentrations indicated.
