## Supplementary Figure S3 for "Assembly reactions of SARS-CoV-2 nucleocapsid protein with nucleic acid"

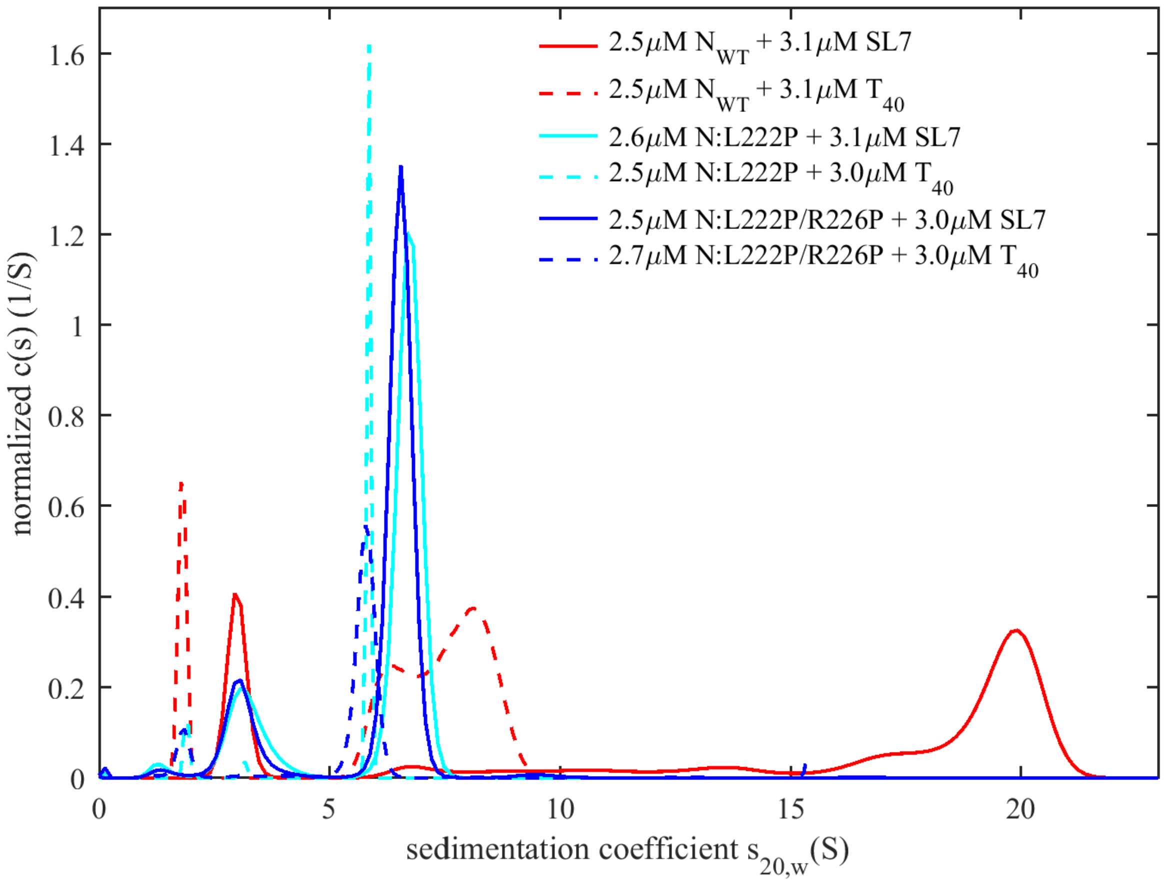


**Supplementary Figure S3: Comparison of RNP formation with oligonucleotide $T_{40}$ and stem-loop SL7.** Shown are sedimentation coefficient distributions recorded at 260 nm for $T_{40}$ (dashed lines) and SL7 (solid lines) with $N_{WT}$ (red) and the LRS mutants N:L222P (cyan) and N:L222P/R226P (blue) all at the same concentrations. Buffer conditions are at moderate ionic strength, identical to those of the RNP assay in MP ($B_{65K}$). This figure reproduces the data from **Supplementary Figure S2** and **Figure 5** to enable a direct visual comparison.
