## Supplementary Figure S4 for "Assembly reactions of SARS-CoV-2 nucleocapsid protein with nucleic acid"

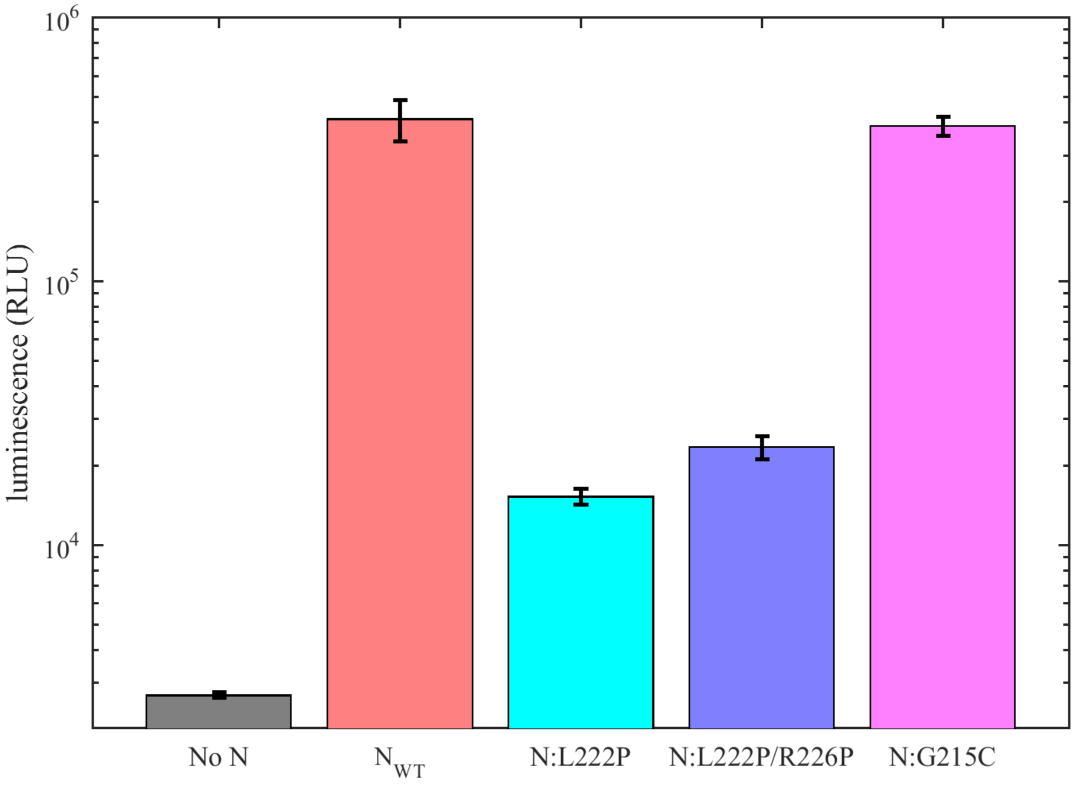


**Supplemental Figure S4: VLP assembly in the presence of GSK3 inhibitor CHIR98014.** Luminescence after incubation of VLP producer cells with 1.25 µM CHIR98014 inhibiting phosphorylation, thereby reducing non-assembly competent populations of intracellular N-protein. The efficiency of VLP formation was measured for different (or lacking) N-protein species indicated and quantified in relative luminescence units.
