## Supplementary Materials and Methods for "Assembly reactions of SARS-CoV-2 nucleocapsid protein with nucleic acid"

*Protein expression and purification*

Wildtype and mutant SARS-CoV-2 N-proteins were expressed and purified as described previously (Zhao et al. 2022; Zhao et al. 2023). Briefly, proteins including N-terminal 6His with TEV cleavage site was cloned into the pET-29a(+) expression vector and transformed into One Shot BL21(DE3)pLysS E. coli (Thermo Fisher Scientific, Carlsbad, CA). After the cells were lysed, the protein was bound to by Ni-NTA column, followed by purification with unfolded and refolded steps to remove residual protein-bound bacterial nucleic acid (Carlson et al. 2020). After $Ni^{2+}$ affinity chromatography, the eluate was subject to cleavage of the 6xHis tag by TEV and purification by affinity and size exclusion chromatography. 95% purity of the protein was confirmed by SDS-PAGE. The ratio of absorbance at 260 nm and 280 nm of ~0.50-0.55 confirmed absence of nucleic acid. The purified protein was dialyzed in the working buffers, either $B_{150Na}$ (20 mM HEPES, 150 mM NaCl, pH. 7.4), $B_{10Na}$ (10 mM NaCl, 10.1 mM $Na_2PO_4$ , 1.8mM $KH_2PO_4$ , 2.7 mM KCl, pH 7.4), or $B_{65K}$ containing 64.8 mM KCl, 5.6 mM NaCl, 24.6 mM HEPES, pH 7.50. The protein concentration was determined by UV-Vis spectrophotometer using $\varepsilon_{280}$ of 43890 $M^{-1}cm^{-1}$.

SARS-CoV-2 NTD including 6His with its sequence, MHHHHHHENLYFQSGSRPQGLPNNTASWFTALTQHGKEDLKFPRGQGVPINTNSSPDDQIGYYRRATRRIRGGDGK MKDLSPRWYFYYLGTGPEAGLPYGANKDGIIWVATEGALNTPKDHIGTRNPANNAAIVLQLPQGTTLPKGFYAE, was cloned into kanamycin-resistant, NdeI/XhoI-digested plasmid pET29a vector (GenScript) in LB with 50mg/ml Kanamycin and 0.5ml IPTG stock (1M). The cells were harvested by centrifugation for 30 minutes at 4000 rpm, and resuspended in lysis buffer (50mM Tris-HCl, pH 7.4, 500mM NaCl, one tablet of protease inhibitor (Sigma S8830), 0.25mg/ml Lysozyme). After sonication at a duty cycle of 50% and an amplitude of 5 for 10 minutes (Branson sonifier model 250 with Branson tapered microtip, 3 mm diameter), the cell debris was pelleted by centrifugation at 35 krpm for 1 hour at 4 °C and the supernatant was then collected for affinity-based purification on a Ni-NTA column (Qiagen). His6-tagged NTD proteins were eluted using elution buffer (20mM Tris, 500mM NaCl, 400mM imidazole at pH 7.4).

Constructs for the SARS-CoV-2 N-protein CTD (247-364) are described in (Karkanitsa et al. 2023). Briefly, an N-terminal His6 tagged N-CTD construct containing a TEV protease cleavage site for tag removal was expressed in 2 liters of E. coli using Dynamite Broth as described in (Taylor et al. 2017) . Cell pellets were lysed by microfluidization, clarified, and subjected to immobilized metal affinity chromatography (IMAC), followed by digestion with TEV protease to remove the purification tag. Samples were then reloaded on an IMAC column, eluted with a shallow gradient from 0-50 mM imidazole, and subjected to size exclusion chromatography on a Superdex S-75 26/60 column (Cytiva) equilibrated in 20 mM HEPES, pH 7.3, 150 mM NaCl, 1 mM TCEP. Purified fractions were analyzed by SDS-PAGE gel and positive fractions were pooled and concentrated for final protein. Final protein was characterized by electrospray ionization mass spectrometry to confirm the appropriate molecular mass.

For all protein constructs, concentrations was determined by UV-Vis spectrophotometer using $\varepsilon_{280}$ of 43890 $M^{-1}cm^{-1}$ (FL-N), 26930 $M^{-1}cm^{-1}$ (NTD) and 16960 $M^{-1}cm^{-1}$ (CTD).

The oligonucleotide $T_{40}$ and stem-loop RNA SL7 were purchased from Integrated DNA Technologies (Skokie, IL), as purified by HPLC and lyophilized. The nucleic acids were reconstituted in buffer $B_{150Na}$. To obtain optimal RNA secondary structure, SL7 was subject to thermal denaturation at 95 °C for 2 min followed by slowly cooling to room temperature over 1-2 hrs. The concentration of the nucleic acids stocks was determined by UV-Vis spectrophotometer using $\varepsilon_{260}$ of 324,600 $M^{-1}cm^{-1}$ ($T_{40}$) and 457,900 $M^{-1}cm^{-1}$ (SL7).

The LRS peptide (N:210-246) was purchased from ABI Scientific (Sterling, VA), purified by HPLC, examined by matrix-assisted laser desorption/ionization for purity and identity, and lyophilized.

*Virus-like particle assay*

To generate virus-like particles with different N mutations, HEK 293T cells were plated at 50,000 cells/well in 150µL of DMEM containing FBS and penicillin/streptomycin in 96 well plates. The next day, 1.5 µg of total plasmids with N (0.33), T20 (0.5), Sv4 (0.0033), and MIEv3 (0.165) at indicated mass ratios were diluted in 75 µL at room temperature with Opti-MEM. Then 4.5 µL of 1% polyethylenimine (pH 7.5) was diluted in 75 µL Opti-MEM; this was used to resuspend plasmid dilutions, followed by incubation at room temperature for 20 minutes. For transfection in 96-well plates, 18.75 µL of resulting transfection mix was added to each well and mixed by pipetting. 48 hours after transfection, 65µL of supernatant was filtered using Pall 0.45 µm AcroPrepTM Advance Plate, Short Tip filter plates (polyethersulfone) by centrifugation (100g for 2 minutes). For VLP infection, 50,000 293T-ACE2/TMPRSS2 cells were added to the VLP-containing filtrate and incubated for 24 hours. For luminescence measurement a Promega Luciferase Assay System (Promega E1501) was used. Briefly, the supernatant was removed, and 20 µL of Promega passive lysis buffer was added to the cells and incubated on a shaker for 20 minutes. Using a Tecan Spark machine, 30 µL of luciferase assay buffer was dispensed into each well and mixed for 60 seconds by shaking. Luminescence was measured using the auto attenuation function and 1000 msec integration time.

For N-protein mutagenesis, plasmids containing B.1 N (Addgene Plasmid #177937) were mutated with primers containing altered codons for their respective mutations:

| | |
|---|---|
| L222P-R226P-Forward | cgccgcactcgctctg**CCC**ctcctggac**CCC**ctcaaccaacttgaatc |
| L222P-R226P-Reverse | gattcaagttggttgag**GGG**gtccaggag**GGG**cagagcgagtgcggcg |
| G215C-Forward | cggcgcggatggcgggcaacggc**TGC**gacgccgcactcgctctg |
| G215C-Reverse | cagagcgagtgcggcgtc**ACG**gccgttgcccgccatccgcgccg |
| L222P-Forward | gacgccgcactcgctctg**CCC**ctcctggaccggctcaacc |
| L222P-Reverse | ggttgagccggtccaggag**GGG**cagagcgagtgcggcgtc |
| R226P-Forward | cgctctgttgctcctggac**CCC**ctcaaccaacttgaatcc |
| R226P-Reverse | ggattcaagttggttgag**GGG**gtccaggagcaacagagcg |

In conditions requiring CHIR98014(CHIR), drug was added immediately after transfection. Media was removed 24 hours later, and 150 µL complete DMEM media was added.

**References**

Carlson CR, Asfaha JB, Ghent CM, Howard CJ, Hartooni N, Safari M, Frankel AD, Morgan DO. 2020. Phosphoregulation of Phase Separation by the SARS-CoV-2 N Protein Suggests a Biophysical Basis for its Dual Functions. *Mol. Cell* 80:1092-1103.e4.

Karkanitsa M, Li Y, Valenti S, Spathies J, Kelly S, Mehalko J, Drew M, Denson J, Putman Z, Fathi P. 2023. Dynamics of SARS-CoV-2 Seroprevalence in a Large US population Over a Period of 12 Months. *medRxiv*.

Taylor T, Denson J-P, Esposito D. 2017. Optimizing Expression and Solubility of Proteins in E. coli Using Modified Media and Induction Parameters. *Methods Mol. Biol.* 1586:65–82.

Zhao H, Nguyen A, Wu D, Li Y, Hassan SA, Chen J, Shroff H, Piszczek G, Schuck P. 2022. Plasticity in structure and assembly of SARS-CoV-2 nucleocapsid protein.Wilson I, editor. *PNAS Nexus* 1:pgac049.

Zhao H, Wu D, Hassan SA, Nguyen A, Chen J, Piszczek G, Schuck P. 2023. A conserved oligomerization domain in the disordered linker of coronavirus nucleocapsid proteins. *Sci. Adv.* 9:eadg6473.
