## Supplementary Figure S1 for "Assembly reactions of SARS-CoV-2 nucleocapsid protein with nucleic acid"

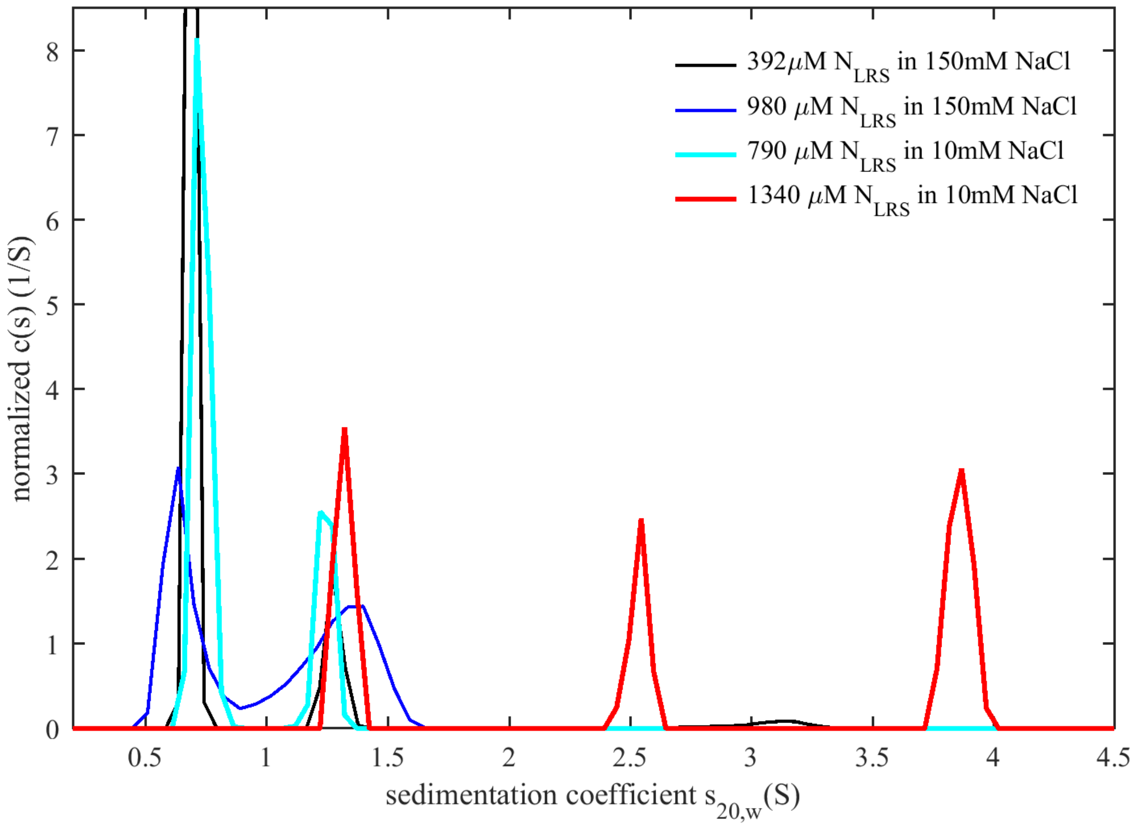


**Supplementary Figure S1: Enhancement of protein-protein interaction in the LRS at low ionic strength.** Shown are sedimentation coefficient distributions recorded for the peptide N:210-246 comprising the LRS transient helix in either phosphate buffered saline with 150 mM NaCl (black and blue curves) or in buffer $B_{10Na}$ containing 10 mM NaCl (cyan and red curves).
